# Flotillin-2 regulates EGFR activation, degradation, and cancer growth

**DOI:** 10.1101/2022.03.11.483779

**Authors:** David J. Wisniewski, Mariya S. Liyasova, Soumya Korrapati, Xu Zhang, Shashikala Ratnayake, Qingrong Chen, Samuel F. Gilbert, Alexis Catalano, Donna Voeller, Daoud Meerzaman, Udayan Guha, Natalie Porat-Shliom, Christina M. Annunziata, Stanley Lipkowitz

**Author notes:** These authors contributed equally to this work (David J. Wisniewski, Mariya S. Liyasova). Corresponding Author: Stanley Lipkowitz, Women’s Malignancies Branch, Center for Cancer Research, National Cancer Institute, National Institutes of Health, Bldg. 10, Room 4B54, Bethesda, MD 20892, USA.

## Abstract

Epidermal growth factor receptor (EGFR) signaling is frequently dysregulated in various cancers. The ubiquitin ligase Cbl (Casitas B-lineage lymphoma proto-oncogene) regulates degradation of activated EGFR through ubiquitination and acts as an adaptor to recruit proteins required for trafficking. We used Stable Isotope Labeling with Amino Acids in Cell Culture (SILAC) mass spectrometry (MS) to compare Cbl complexes with or without epidermal growth factor (EGF) stimulation. We identified over a hundred novel Cbl interactors, and a secondary siRNA screen found that knockdown of Flotillin-2 (FLOT2) led to increased phosphorylation and degradation of EGFR upon EGF stimulation in HeLa cells. In PC9 and H441 cells, FLOT2 knockdown increased EGF-stimulated EGFR phosphorylation, ubiquitination, and downstream signaling, reversible by the EGFR inhibitor erlotinib. CRISPR knockout (KO) of FLOT2 in HeLa cells confirmed EGFR downregulation, increased signaling, and increased dimerization and trafficking to the early endosome. FLOT2 interacted with both Cbl and EGFR. EGFR downregulation upon FLOT2 loss was Cbl-dependent, as co-knockdown of Cbl and Cbl-b restored EGFR levels. Overexpression of FLOT2 decreased EGFR sjgnaling and growth. Overexpression of wild type (WT) FLOT2, but not the soluble G2A FLOT2 mutant, inhibited EGFR phosphorylation upon EGF stimulation in HEK293T cells. FLOT2 loss induced EGFR-dependent proliferation and anchorage-independent growth. Lastly, FLOT2 KO increased tumor formation and tumor volume in nude mice and NSG mice, respectively. These data demonstrated that FLOT2 negatively regulated EGFR activation and dimerization, as well as its subsequent ubiquitination, endosomal trafficking, and degradation, leading to reduced proliferation *in vitro* and *in vivo*.

## Introduction

EGFR is one of the major receptor tyrosine kinases (RTK) in epithelial cells (1). Activation by EGF and other ligands leads to receptor dimerization and phosphorylation at specific tyrosine residues. Activated EGFR activates the RAS/RAF/MAPK and PI3K/AKT pathways, regulating multiple cell functions including proliferation, migration, angiogenesis, and apoptosis (1). EGFR signaling is dysregulated by gene amplification or activating mutations of the receptor in a variety of human malignancies (*e.g.*, lung cancer and glioblastoma) (2, 3). Various targeted therapies have been developed to inhibit EGFR signaling in cancer (4).

The level of activated EGFR in cells is controlled by the Cbl family of ubiquitin ligases (E3), which are recruited to and ubiquitinate the activated receptor leading to lysosomal degradation (5). Loss of Cbl E3 function has been implicated in malignant transformation both in murine models and in human cancers due to hyperactivity of tyrosine kinase-driven pathways (6, 7). Cbl proteins contain a catalytic Ring Finger (RF) responsible for ubiquitination of activated EGFR, surrounded by motifs that interact with multiple proteins via phosphorylated tyrosines, SH2, or SH3 mediated interactions (8). Considering the importance of both Cbl and EGFR in oncogenic transformation, it is critical to gain further understanding of how these pathways regulate oncogenic signaling.

The structure of the N- and C-termini of Cbl allows it to act as an adaptor, recruiting additional proteins required for EGFR internalization and trafficking (8). The effects of Cbl as an adaptor molecule are relatively unknown, and determination of proteins which interact with Cbl as an adaptor may uncover other proteins associated with oncogenic signaling. We thus hypothesized that characterization of proteins that interact with Cbl with and without EGF stimulation will uncover additional mechanisms of EGFR regulation and provide potential therapeutic targets.

In this work, we took two different approaches to study the proteins involved in EGFR activation and trafficking. First, we performed a two-state SILAC MS screen to identify proteins recruited to Cbl with or without EGFR activation. Second, we employed an in-cell ELISA screen to investigate the effect of depletion of specific proteins identified in the SILAC screen on EGFR activation and/or degradation. This approach identified several Cbl interacting proteins for further analysis, including Flotillin-2 (FLOT2).

Flotillin-1 (FLOT1) and FLOT2 are highly conserved and expressed in a variety of cell types (9, 10). The proteins play roles in endocytosis, actin remodeling, signal transduction, and carcinogenesis (11, 12). The two proteins form homo- and hetero-oligomers (13, 14), are membrane associated through myristoylation and palmitoylation, and are used as markers of lipid-raft domains of the plasma membrane (10, 15). Upon EGFR activation, FLOT1 and FLOT2 oligomers increase in size and translocate to the endosomal compartments (13, 16). Flotillins are phosphorylated by the Src family kinase Fyn at Y160 and Y163, which is crucial for their trafficking (17). FLOT1 has been proposed to positively regulate EGFR activation and signaling by regulating EGFR clustering at the plasma membrane and scaffolding with MAPK pathway proteins (11). FLOT2 was also found to be a positive regulator of EGFR phosphorylation in A431 cells (18). Studies linking FLOT2 to EGFR signaling have shown mixed results, as FLOT2 knockdown (KD) has resulted in MAPK activation (19) or MAPK inactivation (20). In the present study we investigated the role of the Flotillin proteins in EGFR activation, signaling, down regulation, trafficking, and growth. Overall, we found that FLOT2 negatively regulates wild type EGFR activation and downstream signaling, downregulation by Cbl-mediated ubiquitination, endocytic trafficking, and cancer cell growth *in vitro* and *in vivo*.

## Results

### SILAC MS identifies Cbl-associated proteins upon EGF stimulation

To identify proteins that interact with Cbl and are modulated upon EGF stimulation, we investigated protein complexes formed with Cbl using SILAC MS. HeLa cell proteins were labeled to compare complexes formed on Cbl without stimulation (L) and upon EGF stimulation (H) (Fig. 1*A*). Following labeling and treatment of HeLa cells, Cbl was immunoprecipitated (IP) from each treatment group and analysis of tryptic peptides was carried out on high-resolution LC-MS/MS Orbitrap mass spectrometer, identifying over 1100 proteins (Table S1). A SILAC ratio cut-off of 1.5 and 0.67 was used for significantly increased and decreased Cbl association, respectively. The interactors of Cbl enriched or depleted upon EGF treatment (H/L > 1.5 or <0.67) are listed in Table 1, with the proteins being studied in a secondary siRNA screen shown in bold.

**Figure 1.**
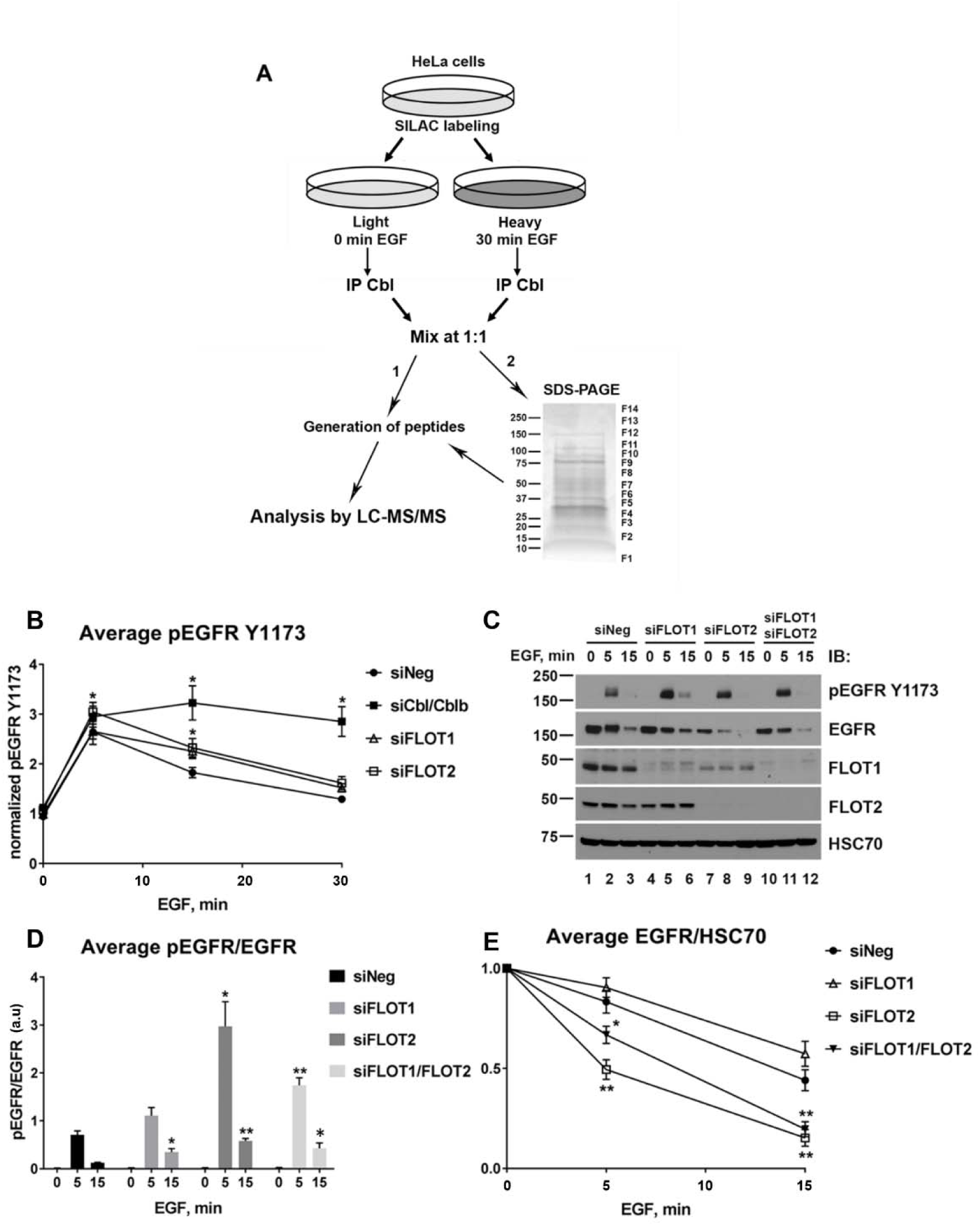
Identification of Cbl interactors with and without EGF stimulation using SILAC MS. **(A)** SILAC MS scheme shows that HeLa cell proteins were labeled with stable isotopes to compare Cbl complexes without EGF (light, L) and with 30 min of 100 ng/mL EGF stimulation (heavy, H). Cbl was immunoprecipitated (IP) from differentially treated cell lysates and the IPs were mixed at 1:1 ratio. The mixed IP was either directly used to generate peptides for MS (1) or was separated by SDS-PAGE (2), divided into fractions as indicated and each fraction was analyzed separately with MS. (B) HeLa cells were transfected with control siRNA (siNeg) or siRNA targeting Cbl/Cbl-b (positive control), FLOT1 or FLOT2 and seeded on 96-well plate. Phospho-EGFR ELISA was used to detect the levels of EGFR phosphorylation at Y1173 upon 100 ng/mL EGF stimulation for indicated time periods. The pEGFR Y1173 signal was normalized to the Janus-green whole-cell staining signal and the average values for three experiments (± SEM) were plotted versus EGF stimulation time. (C) HeLa cells were transfected with control siRNA (siNeg) or siRNA targeting FLOT1, FLOT2 separately and together for 48 hours and then treated with 25 ng/mL EGF for the indicated time periods. The lysates were subjected to western blot analysis with the indicated antibodies. (D) EGFR phosphorylation upon EGF stimulation is shown as an average of pEGFR/EGFR ± SEM calculated by densitometry analysis of western blot (as shown in D) for three independent experiments, with a.u indicating arbitrary units. (E) Degradation of EGFR upon EGF stimulation was calculated by densitometry analysis of western blot in D from seven independent experiments and plotted as an average of EGFR/HSC70 ± SEM. EGFR levels in untreated samples (0 EGF) were set as 1, with a.u indicating arbitrary units.. Asterisk (*) denotes p < 0.05, (**) indicates p < 0.01 as compared to the corresponding time points for siNeg using Student’s *t*-test. MW in kDa is shown to the left of the western blot panels.

**Table 1.**
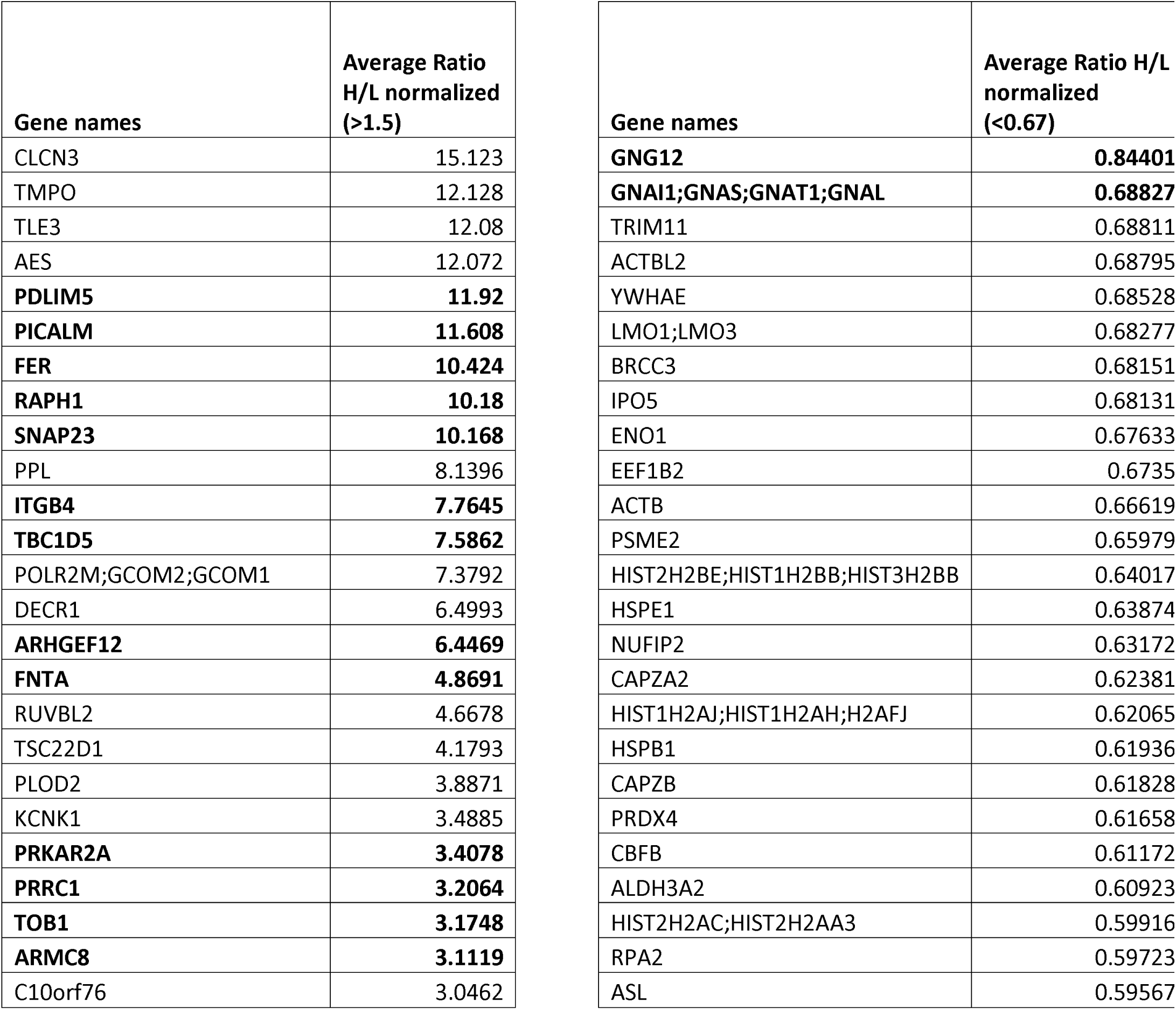
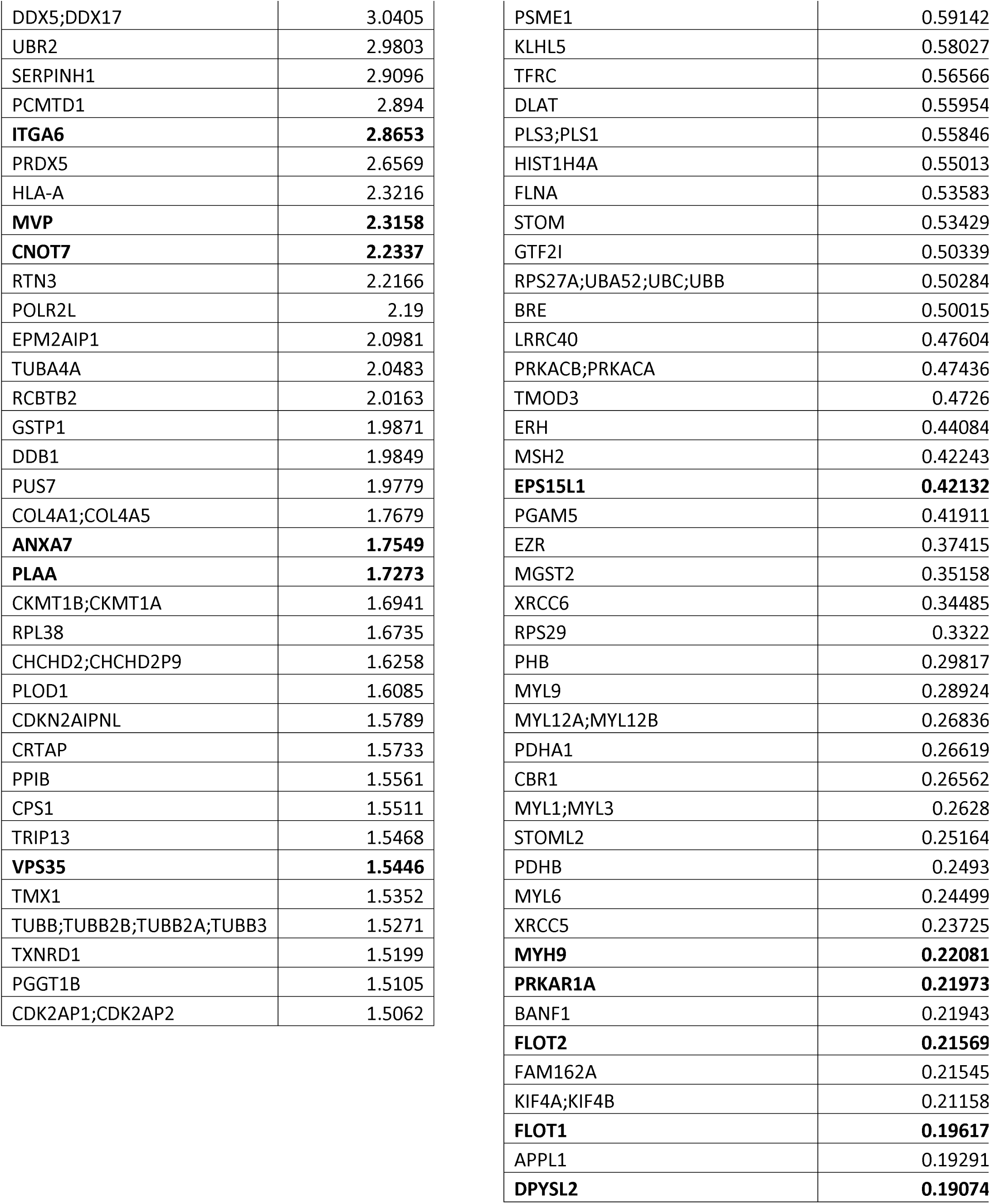
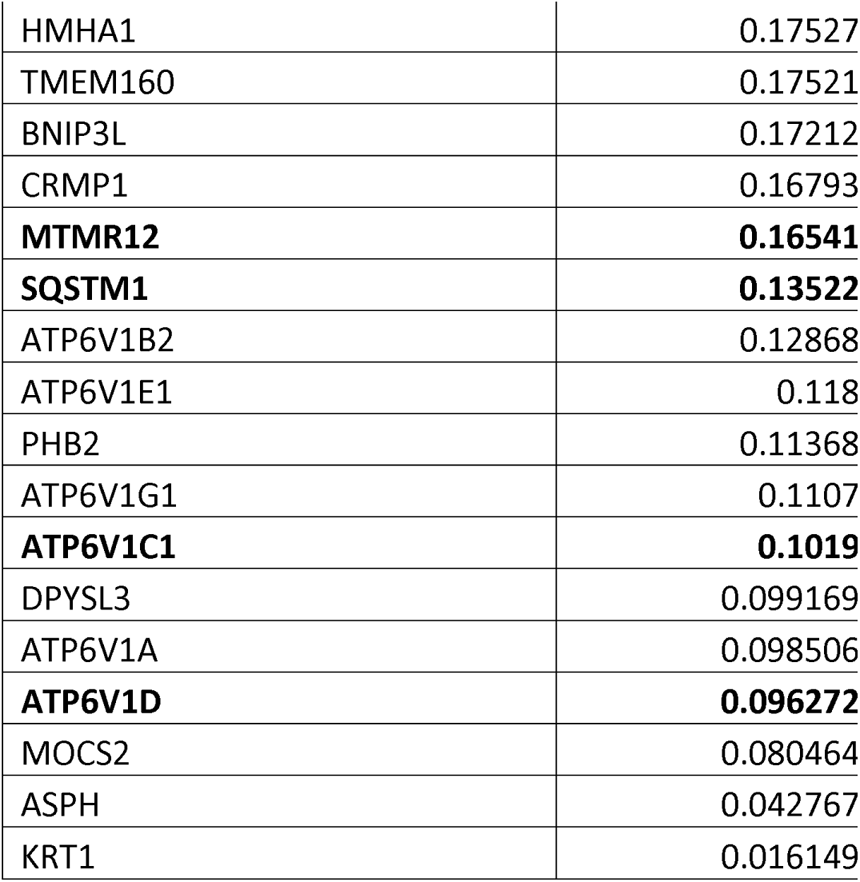
Proteins identified with SILAC MS with significant Cbl association (>1.5 or disassociation (<0.67 H/L)

### pEGFR ELISA identifies proteins involved in EGFR activation

A secondary siRNA screen of pEGFR with stimulation was performed to help select candidates for further analysis. pEGFR (pY1173) signal detected by in-cell ELISA peaked when cells were stimulated with 100 ng/mL EGF at 5 minutes and then gradually decreased to the baseline level by 30 minutes in the presence of EGF (siNeg, Figs. 1*B* and S1*A*). The decline in pEGFR signal with continuous EGF stimulation can be due to EGFR dephosphorylation as well as degradation. We hypothesized that KD of a protein involved in either EGFR phosphorylation, dephosphorylation or degradation would lead to deviation from the pEGFR signal pattern observed with non-targeting siRNA (siNeg). KD of Cbl and Cbl-b served as a positive control as it led to the accumulation of activated EGFR and the persistence of pEGFR signal detected by in-cell ELISA and western blotting (Figs. 1*B*, S1*A* and S1*B*).

Selection of 35 secondary screen target proteins that potentially regulate EGFR trafficking was based on literature reviews and our SILAC MS (Table 2). KD of FLOT1, FLOT2 and SQSTM1/p62 led to the significant increase in pEGFR signal in at least one time point of EGF stimulation (Figs. 1*B* and S1*C*), while the KD of GNG12 decreased pEGFR signal compared to siNeg (Fig. S1*C*). We evaluated the effects of KD of SQSTM1 and GNG12 by western blot and found that the loss of SQSTM1 resulted in increased EGFR phosphorylation while the loss of GNG12 accelerated EGFR degradation (Fig. S1*D*). The KD of GNG12 was confirmed by quantitative reverse transcriptase-polymerase chain reaction (qRT-PCR, data not shown). We chose to move forward with FLOT1/FLOT2 due to their effect on EGFR phosphorylation and degradation and hypothesized that the presence of Flotillin’s at the cell membrane may play a role in EGFR activation at the cell membrane.

**Table 2.** Proteins identified with SILAC MS tested in secondary siRNA screen for their effect on EGFR phosphorylation at Y1173 NS – not significant

| Gene Symbol | Full Gene Name | Gene ID | SILAC H/L | siRNA ID | Tested | Effect on pEGFR |
| --- | --- | --- | --- | --- | --- | --- |
| PDLIM5 | PDZ and LIM domain 5 | 10611 | 11.92 | s20833, s20834, s20835 | 3 | NS |
| PICALM | phosphatidylinositol binding clathrin assembly protein | 8301 | 11.61 | s15799, s15800, s15801 | 4 | NS |
| FER | fer (fps/fes related) tyrosine kinase | 2241 | 10.42 | s5109, s5110, s5111 | 3 | NS |
| RAPH1 | Ras association (RalGDS/AF-6) and pleckstrin homology domains 1 | 65059 | 10.18 | s35177, s35178, s35179 | 2 | NS |
| SNAP23 | synaptosomal-associated protein, 23kDa | 8773 | 10.17 | s16708, s16709, s16710 | 3 | NS |
| ITGB4 | integrin, beta 4 | 3691 | 7.76 | s7583, s7584, s7585 | 3 | NS |
| TBC1D5 | TBC1 domain family, member 5 | 9779 | 7.59 | s229655, s229656, s229657 | 2 | NS |
| ARHGEF12 | Rho guanine nucleotide exchange factor (GEF) 12 | 23365 | 6.45 | s23659, s23660, s23661 | 2 | NS |
| FNTA | farnesyltransferase, CAAX box, alpha | 2339 | 4.87 | s171, s172, s173 | 2 | NS |
| PRKAR2A | protein kinase, cAMP-dependent, regulatory, type II, alpha | 5576 | 3.41 | s11086, s11087, s11088 | 2 | NS |
| PRRC1 | proline-rich coiled-coil 1 | 133619 | 3.21 | s43795, s43796, s43797 | 3 | NS |
| TOB1 | transducer of ERBB2, 1 | 10140 | 3.17 | s19739, s19740, s19741 | 3 | NS |
| ARMC8 | armadillo repeat containing 8 | 25852 | 3.11 | s24623, s24624, s24625 | 3 | NS |
| ITGA6 | integrin, alpha 6 | 3655 | 2.87 | s7492, s7493, s7494 | 2 | NS |
| MVP | major vault protein | 9961 | 2.32 | s19344, s19345, s19346 | 2 | NS |
| CNOT7 | CCR4-NOT transcription complex, subunit 7 | 29883 | 2.23 | s26637, s26638, s26639 | 3 | NS |
| ANXA7 | annexin A7 | 310 | 1.75 | s1398, s1399, s1400 | 2 | NS |
| PLAA | phospholipase A2-activating protein | 9373 | 1.73 | s17934, s17935, s17936 | 3 | NS |
| VPS35 | vacuolar protein sorting 35 homolog (S. cerevisiae) | 55737 | 1.54 | s31374, s31375, s31376 | 4 | NS |
| PRKACB | protein kinase, cAMP-dependent, catalytic, beta | 5567 | 0.47 | s11068, s11069, s11070 | 4 | NS |
| FAM162A | family with sequence similarity 162, member A | 26355 | 0.22 | s25424, s25425, s25426 | 2 | NS |
| MYH9 | myosin, heavy chain 9, non-muscle | 4627 | 0.22 | s222, s223, s224 | 2 | NS |
| ATP6V1D | ATPase, H<sup>+</sup> transporting, lysosomal 34kDa, V1 subunit D | 51382 | 0.1 | s28060, s28061, s28062 | 1 | NS |
| FLOT1 | flotillin 1 | 10211 | 0.2 | s19913, s19914, s19915 | 4 | increased |
| SQSTM1 | sequestosome 1 | 8878 | 0.14 | s16960, s16961, s16962 | 4 | increased |
| GNAI1 | guanine nucleotide binding protein (G protein), alpha inhibiting activity polypeptide 1 | 2770 | 0.69 | s5871, s5872, s5873 | 1 | NS |
| GNG12 | guanine nucleotide binding protein (G protein), gamma 12 | 55970 | 0.84 | s31823, s31824, s31825 | 4 | decreased |
| DPYSL2 | dihydropyrimidinase-like 2 | 1808 | 0.19 | s4269, s4270, s4271 | 2 | NS |
| ATP6V1C1 | ATPase, H<sup>+</sup> transporting, lysosomal 42kDa, V1 subunit C1 | 528 | 0.1 | s1800, s1801, s1802 | 4 | decreased |
| CNPY3 | canopy FGF signaling regulator 3 | 10695 | 0.74 | s21032, s21033, s21034 | 1 | NS |
| PRKAR1A | protein kinase, cAMP-dependent, regulatory, type I, alpha (tissue specific extinguisher | 5573 | 0.22 | s286, s287, s288 | 1 | NS |
|  | 1) |  |  |  |  |  |
| MTMR12 | myotubularin related protein 12 | 54545 | 0.17 | s29177, s29178, s29179 | 2 | NS |
| FLOT2 | flotillin 2 | 2319 | 0.22 | s5284, s5285, s5286 | 4 | increased |
| EPS15L1 | epidermal growth factor receptor pathway substrate 15-like 1 | 58513 | 0.42 | s33903, s33904, s33905 | 1 | NS |

### Knockdown of FLOT2 in HeLa cells increases EGFR phosphorylation and degradation

We hypothesized that Flotillin’s may regulate EGFR activation or trafficking. To test this, FLOT1 and FLOT2 were KD in HeLa cells either alone or in combination and EGFR phosphorylation was assessed upon EGF stimulation. The KD of FLOT2 also decreased the levels of FLOT1, while FLOT1 KD had no significant effect on the levels of FLOT2 protein (Figs. 1*C* and S2*A*). The loss of FLOT1 with FLOT2 KD was confirmed by three separate FLOT2 siRNAs in both HeLa and H441 cells (Fig. S2, *B* and *C,* respectively). The loss of FLOT1 when FLOT2 is KD is consistent with previous work showing that the stability of FLOT1 is dependent on the presence of FLOT2 (14). Further, CRISPR-mediated FLOT2 knockout (KO) also reduced FLOT1 levels in HeLa cells (Fig. S2*D*).

Stimulation of HeLa cells with EGF induced EGFR phosphorylation at Y1173 and EGFR degradation. To quantify the effect of KD of FLOT1 or FLOT2 on EGFR phosphorylation, the intensities of pEGFR pY1173 were normalized to the intensities of EGFR bands (Fig. 1, *C* and *D*). KD of FLOT1 resulted in a slight increase in EGFR phosphorylation upon EGF stimulation compared to the negative control; however, this effect was statistically significant only at the 15 minute time point. In contrast, KD of FLOT2 led to a decrease in steady-state levels of EGFR as well as an increase in EGF-induced EGFR phosphorylation at Y1173 at 5 and 15 minutes (Fig. 1, *C* and *D*). The KD of both FLOT1 and FLOT2 significantly increased EGFR phosphorylation upon 5 and 15 minutes of EGF stimulation (Fig. 1, *C* and *D*). This effect was not limited to a specific tyrosine residue, because a similar pattern was observed when the phosphorylation at Y845 was analyzed (Fig. S2*A*). Further, increases in EGFR phosphorylation relative to total EGFR and decreases in steady-state levels of EGFR were observed with three separate siFLOT2 constructs in both HeLa and H441 cells (Figs. S2 *B* and *C,* respectively).

To test the effect of the FLOT1 and FLOT2 KD on EGFR degradation, the intensities of EGFR signal on western blot were normalized to the internal control (HSC70). The resulting ratios were plotted versus time of EGF stimulation (Fig. 1*E*). The KD of FLOT2 as well as the combined KD of FLOT1 and FLOT2 led to significant increases in EGFR degradation. Since FLOT2 KD showed the strongest effects on EGFR phosphorylation and down regulation, hereafter we investigated the role of FLOT2 on EGFR activation and down-regulation.

### FLOT2 and Cbl interaction decreases upon EGF stimulated EGFR activation

To confirm the interaction and specificity between FLOT2 and Cbl identified by SILAC, we first tested co-immunoprecipitation (IP) of Cbl with FLOT2 using control and FLOT2 KO HeLa cells. We created FLOT2 KO clones in HeLa cells using the CRISPR/cas9 system (Fig. S3*A*). Both alleles were targeted and resulted in complete protein loss. In parallel, two control clones were created by targeting the *eGFP* gene, normally absent from HeLa cells. We immunoprecipitated FLOT2 in both control and FLOT2 KO HeLa cells and observed a specific co-immunoprecipitation of Cbl in the control cells, not in the FLOT2 KO cells (Fig S3*B*). KO of FLOT2 was confirmed in the whole cell lysate (WCL) and FLOT2 IP. In the SILAC experiment the interaction between Cbl and FLOT2 decreased with EGF stimulation (Table 1 and Table 2). To test this, we stimulated HeLa cells with 100 ng/mL EGF for up to 30 minutes, as done in the SILAC experiment, followed by FLOT2 IP. Again, we found an interaction between Cbl and FLOT2 in unstimulated cells that decreased with EGF activation (Fig S3*C*). Overall, this confirms the interaction between Cbl and FLOT2, and that with EGF stimulation this interaction is decreased.

### Knockdown of FLOT2 in H441 cells increases EGFR phosphorylation, ubiquitination and signaling

Cancer genomics data sets in cBioPortal (www.cbioportal.org) indicated that FLOT2 is amplified in lung adenocarcinoma cell line H441 which, similar to HeLa, has wild type EGFR (21). We confirmed a higher FLOT2 expression than HeLa cells, and EGF treatment similarly induced EGFR activation and ubiquitination in H441 (Fig. S4*A*). FLOT2 KD significantly increased EGF-induced EGFR phosphorylation at multiple tyrosine residues compared to the cells transfected with siNeg (Fig. 2*A,* S4*B*). Similar to HeLa cells, FLOT2 KD in H441 reduced baseline levels of EGFR and FLOT1 (Fig. 2*A*, S2*B*). Moreover, FLOT2 KD led to an increase in phosphorylation of MAPK, a downstream target of EGFR signaling, even without EGF stimulation (Fig. 2*A*, lanes 4-6).

**Figure 2.**
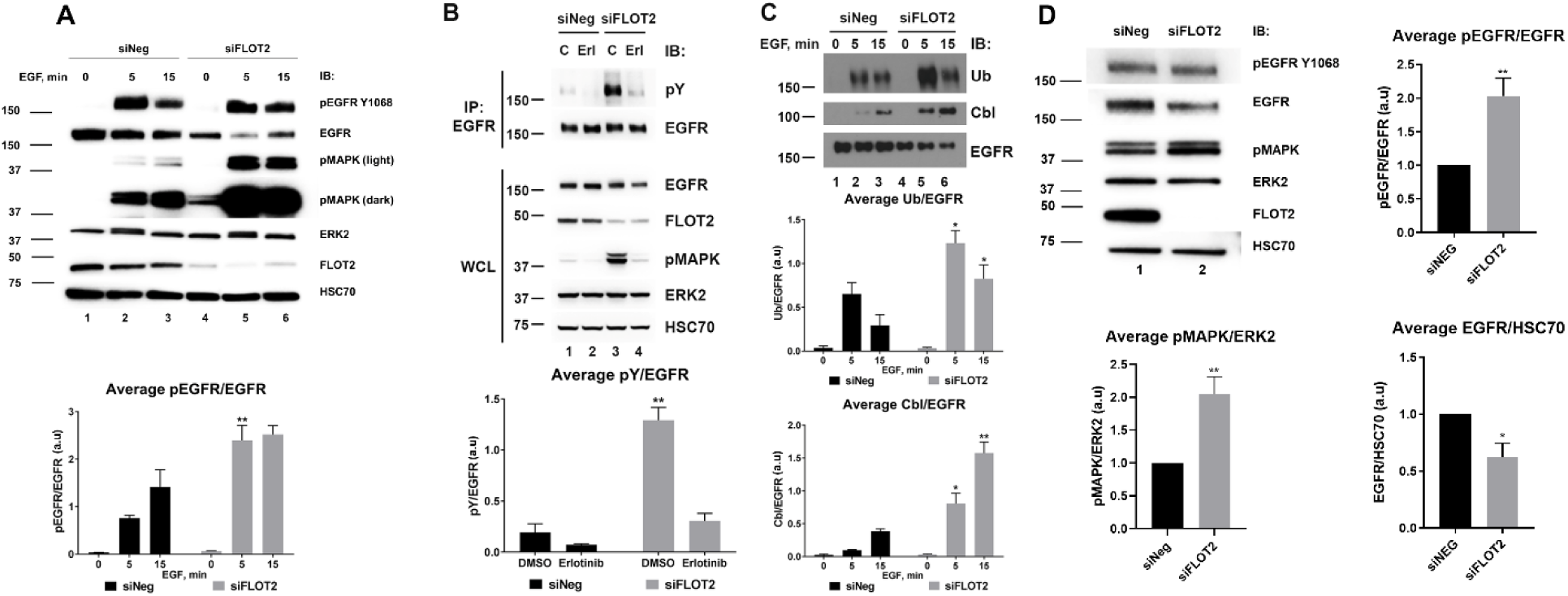
Knockdown of FLOT2 in H441 and PC9 cells increases EGFR phosphorylation and signaling. **(A)** FLOT2 was knocked down in H441 cells by specific siRNA for 72 hours and its effect on EGFR signaling upon stimulation with 100 ng/mL EGF was compared to that of non-targeting siRNA (siNeg). The graph under the western blot panels shows an average of pEGFR/EGFR ± SEM calculated by densitometry analysis of western blot as above for three independent experiments, with a.u indicating arbitrary units. (B) FLOT2 was knocked down in H441 cells as in A, but for the last 24 hours of siRNA transfection, the cells were treated with 10 μM erlotinib. To study the effect on EGFR phosphorylation, EGFR was immunoprecipitated (IP EGFR) and probed to detect tyrosine phosphorylation with anti-pY (4G10) antibodies and EGFR. The graph below the western blot panels shows an average of pY/EGFR ± SEM calculated by densitometry analysis of EGFR IP for three independent experiments, with a.u indicating arbitrary units. Whole cell lysates (WCL) were analyzed with specific antibodies to detect EGFR, FLOT2, pMAPK, ERK2 and HSC70 as indicated. (C) EGFR was immunoprecipitated from lysates in A and analyzed with anti-ubiquitin (UB), anti-Cbl and anti-EGFR ABs. The graphs below the western blot panels show an average of Ub/EGFR and Cbl/EGFR ± SEM calculated by densitometry analysis of corresponding bands of EGFR IP for four and three independent experiments, respectively, with a.u indicating arbitrary units. In all panels, asterisk (*) denotes p < 0.05, (**) indicates p < 0.01 as compared to the corresponding time points for siNeg or DMSO using Student’s *t*-test. MW in kDa is shown to the left of the western blot panels. (D) FLOT2 was knocked down in PC9 cells by specific siRNA for 48 hours and the effect on EGFR and MAPK signaling was compared to non-targeting siRNA (siNeg). The graphs show an average of pEGFR/EGFR ± SEM, pMAPK/ERK2 ± SEM, and EGFR/HSC70 ± SEM calculated by densitometry analysis of western blot for five independent experiments, with a.u indicating arbitrary units. Asterisk (*) denotes p < 0.05, (**) indicates p < 0.01 as compared to siNeg using Student’s *t*-test. MW in kDa is shown to the left of the western blot panels.

We hypothesized that the EGFR is constitutively activated by the KD of FLOT2, resulting in increased basal MAPK phosphorylation. To test this, EGFR was immunoprecipitated and the basal phosphorylation was assessed with anti-phosphotyrosine antibodies. As predicted, KD of FLOT2 resulted in an almost 14-fold increase in basal EGFR phosphorylation as compared to negative control (Fig. 2*B*, top panel, compare lane 3 to lane 1). In addition, the treatment of the cells with the EGFR tyrosine kinase inhibitor erlotinib after FLOT2 KD reduced EGFR phosphorylation levels and pMAPK levels down to baseline (Fig. 2*B*, compare lane 4 to lane 3). Together these data indicate that the increase in pMAPK seen in the siFLOT2 H441 cells is driven by activated EGFR.

EGFR was immunoprecipitated from H441 cells with or without EGF stimulation and probed for ubiquitin and Cbl recruitment (Fig. 2*C*). The KD of FLOT2 resulted in an increase in EGFR ubiquitination and a greater association with Cbl compared to the negative control upon 5 minutes of EGF stimulation. Overall, these data suggest that KD of FLOT2 increases both basal and EGF-induced EGFR phosphorylation, leading to activated downstream signaling, increased association with Cbl, and increased EGFR ubiquitination.

### Knockdown of FLOT2 in PC9 cells induces EGFR and MAPK phosphorylation, and EGFR downregulation

To further confirm the role of FLOT2 in EGFR signaling, we used the lung adenocarcinoma cell line PC9, which has an activating EGFR mutation (deletion E746_A750) (21, 22). These cells exhibit significantly less FLOT2 than both HeLa and H441 cells, and exhibit EGFR-dependent growth, as they are sensitive to erlotinib at low concentrations of 100 and 1000 nM (Fig. S4 *C* and *D*). Indeed, these cells also exhibit increased EGFR and MAPK phosphorylation with FLOT2 KD, as well as downregulation of EGFR (Fig. 2*D*). Considering the significant signaling induction of MAPK in PC9 and H441 cells, observed to be through EGFR activation, we investigated whether other RTK’s contributed to the MAPK activation with FLOT2 KD. In both H441 (Fig. S5*A*) and PC9 (Fig. S5*B*), cabozantinib (VEGFR, MET, KIT, RET, AXL inhibitor) and defactinib (FAK inhibitor) did not reduce the observed pMAPK induction by FLOT2 KD, while erlotinib (EGFR inhibitor) did. There was no statistical effect on MET activation by FLOT2 KD in either cell line. FLOT2 KD also did not affect FAK activation in H441 cells, however FAK activation was decreased with loss of FLOT2 in PC9 cells. Lastly, cabozantinib inhibited pMET and defactinib inhibited pFAK at the concentrations used, confirming that the drugs were active at these concentrations (Fig S5).

### CRISPR knockout of FLOT2 downregulates EGFR levels and increases EGFR phosphorylation

Confirming our siFLOT2 experiments, the basal levels of EGFR were reduced upon CRISPR mediated KO of FLOT2 (Fig. 3*A*, compare lanes 7-12 to lanes 1-6). EGF-induced phosphorylation of EGFR at Y1068 was significantly increased in both FLOT2 KO clones when compared to the C1A control clone and normalized to total EGFR (Fig. 3, *A* and *B*). This effect was also confirmed by EGFR IP and probing with anti-phosphotyrosine antibodies (Fig. 3*A*).

**Figure 3.**
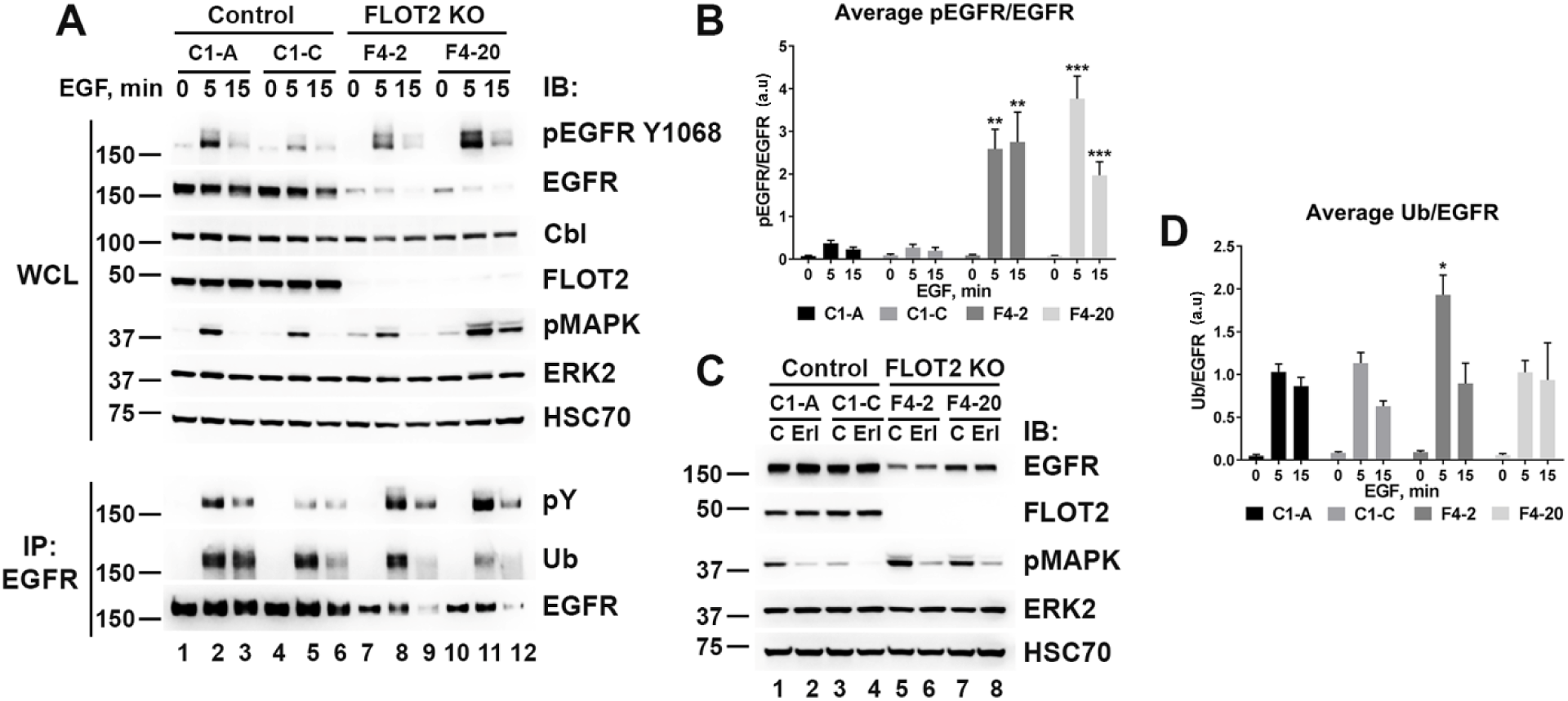
CRISPR knockout of FLOT2 increases EGFR activation and signaling, while decreasing EGFR steady-state levels. **(A)** Two control (C1-A, C1-C) and two FLOT2 KO CRISPR (F4-2, F4-20) clones were created in HeLa cells and whole-cell lysates (WCL, upper panel) were analyzed on western blot with the indicated antibodies. EGFR was immunoprecipitated (IP EGFR, lower panel) and analyzed for phosphorylation (pY 4G10 AB), ubiquitination (Ub) and EGFR levels. (B) The graph shows the average EGFR phosphorylation at Y1068 for five experiments calculated as the intensity of pEGFR to EGFR bands on WCL as in A, with a.u indicating arbitrary units. (C) CRISPR clones were treated with DMSO (C) or 10 μM erlotinib (Erl) for 24 hours. Then the lysates were analyzed on western blot with the antibodies indicated to the right of the panel. **(D)** The graph shows the average EGFR ubiquitination for four experiments calculated as the intensity of Ub to EGFR bands on IP EGFR as in A, with a.u indicating arbitrary units. In all panels, asterisk (*) denotes p < 0.05, (**) indicates p < 0.01 and (***) indicates p < 0.001 as compared to the corresponding time points for C1-A using Student’s *t*-test. MW in kDa is shown to the left of the panels.

Consistent with increased EGFR activation, FLOT2 KO led to an increase in phosphorylation of MAPK without EGF stimulation. This effect was most prominent in the F4-20 clone (Fig. 3*A*, compare lanes 7 and 10 to lanes 1 or 4 of WCL). Similar to H441 cells (Fig. 2*B*), the effect of FLOT2 KO on MAPK activation in the absence of exogenous EGF was blocked by the EGFR inhibitor erlotinib, confirming that the MAPK phosphorylation is being driven by EGFR activity (Fig. 3*C*).

To test whether EGFR phosphorylation led to increased EGFR ubiquitination, EGFR was immunoprecipitated from control or FLOT2 KO clones stimulated with EGF and probed for ubiquitin (Fig. 3*A*, IP EGFR). The two clones had different ubiquitination profiles (Fig. 3*D*). The F4-2 clone had increased EGFR ubiquitination upon 5 mintes of EGF stimulation compared to the control clones, however there was not a discernible difference in ubiquitination in F4-20. In summary, we observed that the depletion of FLOT2 leads to increased EGFR activation and signaling as well as reduced steady-state levels of the receptor. The differential pattern of EGFR ubiquitination in FLOT2 KO clones may reflect different adaptation mechanisms to compensate for the loss of FLOT2 in the stable KO clones.

### FLOT2 prevents EGFR downregulation by Cbl/Cbl-b

As loss of FLOT2 increased ubiquitination and downregulation of EGFR, we investigated whether this downregulation was dependent on Cbl/Cbl-b. Cbl/Cbl-b KD prevented EGF-stimulated EGFR down regulation in the control (C1-A) HeLa cells (Fig. 4*A;* compare lanes 2 and 4), confirming that Cbl and Cbl-b are required for EGF-induced EGFR downregulation, which is well-established by our group and others (23, 24). When Cbl and Cbl-b were knocked down in FLOT2 KO cells, basal EGFR levels increased, confirming that FLOT2 loss leads to Cbl/Cbl-b dependent downregulation of EGFR (Fig. 4*A*; compare lanes 5 and 6 to lanes 7 and 8, and lanes 5 and 7 to 1; and compare bars 1, 5, and 7 in the quantification in Fig. 4*B*). It should be noted that with FLOT2 KO and Cbl/Cbl-b KD and EGF stimulation, there is a slight reduction in total EGFR levels compared to unstimulated cells (compare lane 7 and 8), however this change is not statistically significant. This could be due to incomplete KD of Cbl/Cbl-b or alternatively there may be other Cbl/Cbl-b independent mechanisms of downregulation. Similarly, in H441 cells the loss of EGFR with FLOT2 KD were reversed by siRNA mediated KD of Cbl/Cbl-b (Fig. 4, *C* and *D*; compare lane/bars 1,3 and 4).

**Figure 4.**
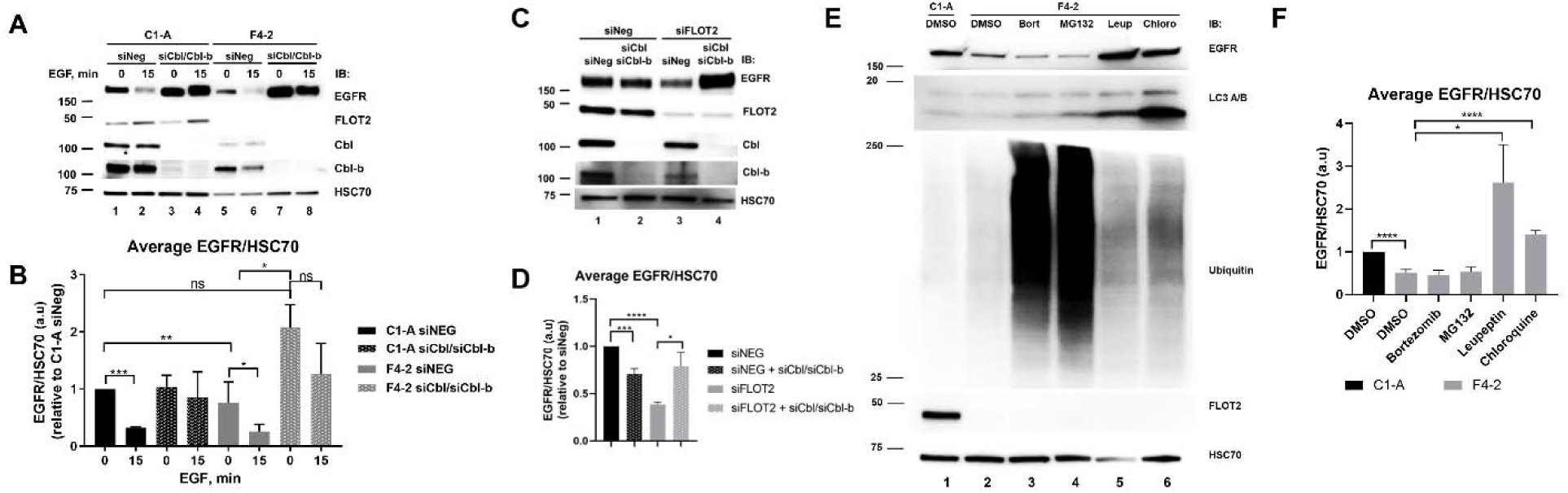
Downregulation of EGFR in response to FLOT2 knockout/knockdown is dependent on Cbl/Cbl-b and lysosomal signaling. **(A)** Cbl and Cbl-b were knocked down in control (C1-A) and FLOT2 CRISPR KO (F4-2) HeLa cells by specific siRNA for 72 hours, and their effect on downregulation of EGFR upon stimulation with 25 ng/mL EGF was compared to that of non-targeting siRNA (siNeg). (B) The average of EGFR/HSC70 ± SEM calculated by densitometry analysis for three independent experiments as shown in A, with a.u indicating arbitrary units.. Asterisks (***) denotes p < 0.001, (**) denotes p <0.01, (*) denotes p < 0.05, and non-significant (ns; p >0.05) using Student’s *t*-test. (C) FLOT2, Cbl and Cbl-b were knocked down in H441 cells by siRNA for 72 hours, and their effect on downregulation of EGFR was compared to that of non-targeting siRNA (siNeg). Western blot analysis with anti-Cbl, Cbl-b, and FLOT2 ABs demonstrate successful knockdown of their targets. (D) The average of EGFR/HSC70 ± SEM calculated by densitometry analysis for three independent experiments as shown in C, with a.u indicating arbitrary units. Asterisks (****) denotes p < 0.0001, (***) denotes p <0.001, and (*) denotes p < 0.05 using Student’s *t*-test. MW in kDa is shown to the left of the panels. (E) Control (C1-A) HeLa cells were treated with DMSO, and FLOT2 KO CRISPR (F4-2) HeLa cells were treated with DMSO, Bortezomib (10 nM), MG132 (5 µ M), Leupeptin (100 µ M) or Chloroquine (25µ M) for 48 hours, lysed and analyzed by western blot. (F) The average EGFR/HSC70 ± SEM calculated by densitometry analysis for three independent experiments as shown in E, with a.u indicating arbitrary units. Asterisks (****) denotes p < 0.0001, and (*) denotes p < 0.05 using Student’s *t*-test. MW in kDa is shown to the left of the panels.

Cbl mediates primarily multiple sites of mono- and di-ubiquitination of the activated EGFR which is a signal for endocytic trafficking to the lysosome for degradation (25). To discern whether the degradation of EGFR seen upon FLOT2 loss was due to the lysosomal or proteasomal pathway, we treated HeLa cells with proteasomal inhibitors bortezomib or MG132, or lysosomal inhibitors leupeptin or chloroquine for 48 hours and observed EGFR levels by western blot (Fig 4*E* and *F*). Lysosomal inhibition was confirmed by increases in LC3 protein expression, and proteasomal inhibition was confirmed by increases in total ubiquitination of whole cell lysate (Fig 4E; lower panels). Both of the lysosomal inhibitors, but not the proteasomal inhibitors, rescued total EGFR levels in FLOT2 KO HeLa cells, indicating that the downregulation of EGFR is due to the lysosomal pathway.

### Loss of FLOT2 increases EGFR dimerization

Since loss of FLOT2 results in increased activation of EGFR, we investigated whether this activation was a result of dimerization of the receptor. Proximity ligation assay (PLA) allows visualization of homodimerization of two EGFR molecules which are separately bound by individual EGFR probes (26). Upon EGF stimulation, EGFR dimerization was observed by increased PLA signal in control HeLa cells when both plus and minus probes are used, while no signal was seen when either probe alone is used, indicating there is no off-target fluorescence in individual probe treatment (Fig. S6, *A* and *B*).

FLOT2 KO HeLa cells exhibited statistically significant increases in EGFR dimerization in the absence of exogenous EGF as evidenced by an increase in green fluorescence using PLA (Fig. 5, *A* and *B*), as did H441 cells upon FLOT2 KD (Fig. 5, *C* and *D*). To investigate whether membrane localized EGFR levels could explain the changes in activation and downregulation, subcellular fractionation was performed in order to determine EGFR levels in the membrane. Membrane localized EGFR also decreased with FLOT2 KO (Fig. S6*C*). EGFR was normalized to calreticulin, a known membrane localized protein whose expression should not change with FLOT2 loss. We investigated the role of membrane rafts on EGFR downregulation and dimerization through the use of simvastatin, a cholesterol synthesis inhibitor. Simvastatin reduced total cholesterol levels in HeLa cells at 24 hours (Fig. S6*D*), however it did not affect downregulation of total EGFR levels with FLOT2 loss (Fig. S6*E*). Further, cholesterol depletion did not reduce the FLOT2 KO induced EGFR dimerization in HeLa cells, instead increasing dimerization of EGFR in FLOT2 KO HeLa cells even more (Fig. S6*F*). Taken together, these data signify that loss of FLOT2 increases EGFR dimerization in both HeLa and H441, EGFR downregulation is consistently observed in the membrane, and the dimerization and downregulation is not due to lipid raft presence or cholesterol synthesis.

**Figure 5.**
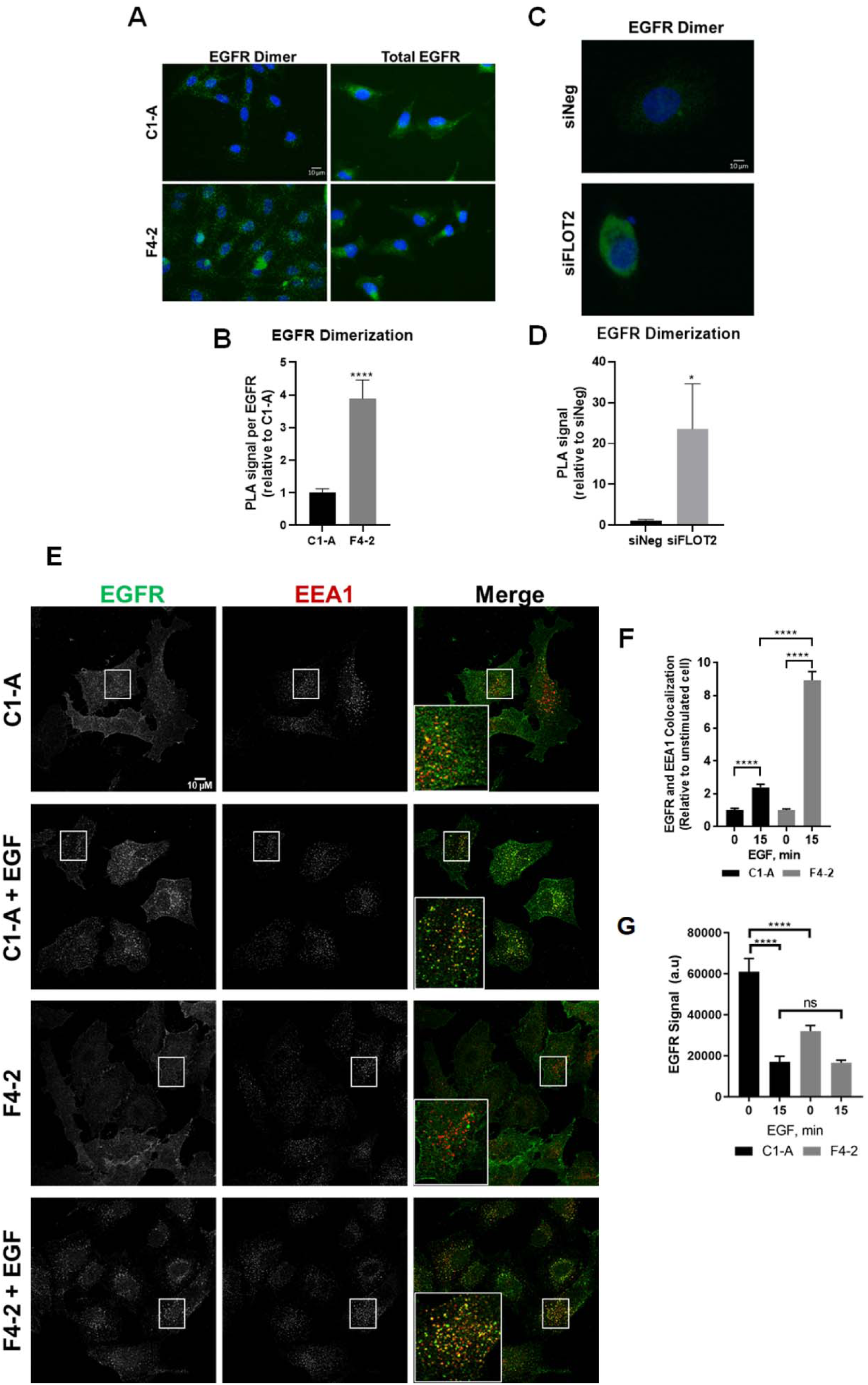
Knockout of FLOT2 significantly increases EGFR dimerization and trafficking to the early endosome upon EGF stimulation. **(A)** Control (C1-A) and FLOT2 KO CRISPR (F4-2) HeLa cells were treated with PLUS and MINUS probes conjugated to EGFR ABs in a proximity ligation assay (PLA). Dimerized EGFR was visualized by green fluorescence, when both PLUS and MINUS probes bound to EGFR were within 40 nm. Nuclei were stained with DAPI. Total EGFR was visualized with un-conjugated EGFR ABs and an Alexa-Fluor 488 secondary AB. Scale bar = 10 microns. (B) Fluorescence was quantified using ImageJ, PLA signal to EGFR was calculated, and normalized to C1-A control. The graph shows the average (± SEM) of three independent experiments. Asterisks (****) denotes p < 0.0001 using Student’s *t*-test. (C) FLOT2 was knocked down in H441 cells by specific siRNA for 72 hours, and the cells were stained with PLUS and MINUS probes and DAPI as in A. Scale bar = 10 microns. (D) Fluorescence was quantified using ImageJ and normalized to siNeg control. The graph shows the average (± SEM) of three independent experiments. Asterisks (*) denotes p < 0.05 using Student’s *t*-test. (E) Control (C1-A) and FLOT2 (F4-2) CRISPR KO HeLa cells were treated with or without 25 ng/mL EGF for 15 minutes, and were subsequently stained with EGFR AB and EEA1 AB, followed by Alexa-Fluor 488 and Alexa-Fluor 594 fluorescent secondary AB, respectively. Insets show higher resolution overlapping EGFR and EEA1 pixels (yellow). Scale bar = 10 microns. (F) Overlapping EGFR and EEA1 pixels were calculated using ImageJ. Stimulated C1-A cells were normalized to unstimulated C1-A cells, and stimulated F4-2 cells were normalized to unstimulated F4-2 cells. The graph shows the average (± SEM) of three independent experiments. Asterisks (****) denotes p < 0.0001 using Student’s *t*-test. (G) Total EGFR signal was quantified using ImageJ, and plotted as arbitrary units (a.u). The graph shows the average (± SEM) of three independent experiments. Asterisks (****) denotes p < 0.0001, and non-significant (ns; p >0.05) using Student’s *t*-test.

### Knockout of FLOT2 in HeLa cells increases EGF-induced trafficking of EGFR to the early endosome

To further investigate the role FLOT2 plays in trafficking of EGFR, we utilized confocal microscopy and examined the co-localization of EGFR with endosomal or lysosomal markers in FLOT2 KO cells. Upon EGF stimulation, EGFR exhibited a statistically significant increase in trafficking to the early endosome in FLOT2 KO HeLa cells compared to control KO cells, as evidenced by a higher fraction of EGFR co-localization with Early Endosome Antigen 1 (EEA1) when normalizing stimulated C1-A to unstimulated C1-A, and stimulated F4-2 to unstimulated F4-2 (Fig. 5, *E* and *F*). This study indicates that loss of FLOT2 enhanced endosomal trafficking in response to EGF stimulation. Confocal imaging visualizing total EGFR confirmed prior observation of EGFR downregulation in unstimulated FLOT2 KO HeLa cells (Fig. 5*G*). Co-localization of EGFR with lysosomal marker LAMP1 was significantly increased in the FLOT2 KO cells with EGF stimulation compared to unstimulated FLOT2 KO cells, while the increase in the control cells was statistically borderline (Fig. S7). Of note, there was no statistically significant change in lysosomal colocalization when comparing EGF stimulated control to FLOT2 KO cells (Fig S7), although we did observe inhibition of the lysosome with chloroquine or leupeptin rescued total EGFR levels (Fig. 4*E* and *F*).

### FLOT2 overexpression decreases EGFR phosphorylation and ubiquitination

We established HeLa cells that stably overexpress FLAG-Myc-FLOT2 (WT), with empty vector as a control (Vector), and confirmed expression by immunoblotting (Fig. 6*A*). FLOT2 overexpression significantly decreased EGF-induced EGFR phosphorylation compared to vector control (Fig. 6*B*).

**Figure 6.**
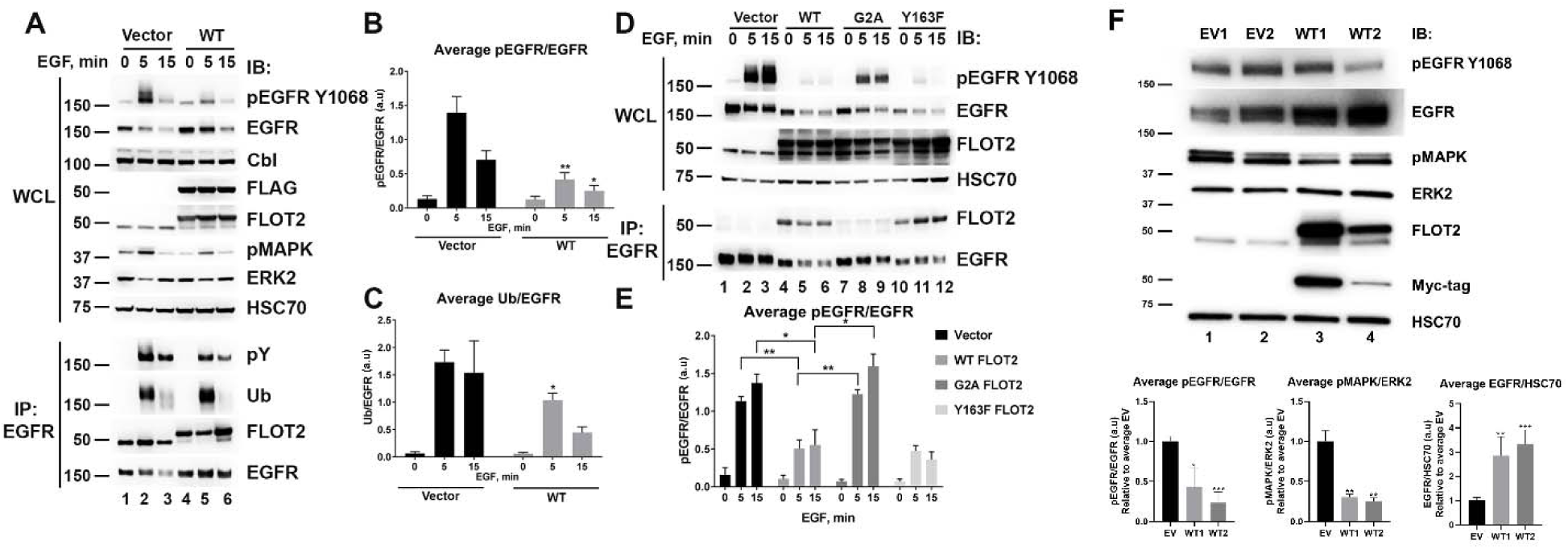
FLOT2 overexpression decreases EGFR phosphorylation and signaling. **(A)** Control (Vector) and FLAG-FLOT2 overexpressing (WT) clones were created in HeLa cells and whole-cell lysates (WCL, upper panel) were analyzed on western blot with the indicated antibodies. EGFR was immunoprecipitated (IP EGFR, lower panel) and analyzed for phosphorylation (pY 4G10 AB), ubiquitination (Ub), interaction with FLOT2 and EGFR levels. (B) The graph shows the average (± SEM) EGFR phosphorylation at Y1068 for six experiments calculated as a ratio of the intensity of pEGFR to EGFR bands on WCL as in A, with a.u indicating arbitrary units. (C) The graph shows the average (± SEM) EGFR ubiquitination for five experiments calculated as the intensity of Ub to EGFR bands on IP EGFR as in A, with a.u indicating arbitrary units. (D) Vector, FLAG-tagged WT FLOT2 and FLOT2 mutants (G2A and Y163F) were overexpressed in HEK293T cells for 48 hours and the cells were stimulated with 100 ng/mL EGF as indicated. The lysates (WCL, upper panel) as well as EGFR IP (lower panel) were analyzed on western blot with the indicated antibodies. (E) The effect of FLOT2 overexpression on EGFR phosphorylation was assessed by normalizing the intensities of pEGFR bands to corresponding EGFR bands of WCL as in D, with a.u indicating arbitrary units. The graph shows the average (± SEM) of three independent experiments. In all panels, asterisk (*) denotes p < 0.05, (**) indicates p < 0.01 as compared to the corresponding time points of Vector (except E) using Student’s *t*-test. MW in kDa is shown to the left of the western blot panels. (F) Empty vector (EV) or overexpressing wild-type FLOT2 (WT) stable PC9 cells were lysed and analyzed by western blot. The graphs show an average of pEGFR/EGFR ± SEM, pMAPK/ERK2 ± SEM, and EGFR/HSC70 ± SEM calculated by densitometry analysis of western blot for three independent experiments, with a.u indicating arbitrary units. Densitometry was normalized to the average signal of EV1 and EV2, and asterisk (*) denotes p < 0.05, (**) indicates p < 0.01, and (***) indicates p<0.001 as compared to empty vector average using Student’s *t*-test. MW in kDa is shown to the left of the western blot panels.

The steady-state levels of EGFR were higher with FLOT2 overexpression (Fig. 6*A,* compare lane 4 to lane 1). There was a 1.7-fold decrease in EGFR ubiquitination in the FLOT2 overexpressing clone compared to the control clone at 5 minutes of EGF stimulation (Fig. 6, *A* and *C*). Endogenous as well as overexpressed FLOT2 co-immunoprecipitated with EGFR irrespective of EGF stimulation, indicating a constitutive interaction between FLOT2 and EGFR proteins (Figs. 6*A* and S8). This finding is in agreement with published data that Flotillins form constitutive complexes with EGFR (11). Thus, overexpression of FLOT2 in HeLa cells reduced EGFR phosphorylation and ubiquitination induced by EGF activation.

### Soluble G2A FLOT2 mutant does not affect EGFR phosphorylation, but WT and Y163F mutant do

We generated two FLOT2 mutants: a G2A mutant, lacking the glycine residue which is myristolyated, making it unable to associate with the plasma membrane (15), and a Y163F FLOT2 mutant that is deficient in phosphorylation by the Src family kinase Fyn (17). Upon overexpression, G2A mutant is fully soluble in the cytosol (15), while Y163F mutant remains fully associated with the plasma membrane (16, 17). These mutants were overexpressed in HEK293T cells along with EGFR and Cbl plasmids and their effect on EGFR phosphorylation was compared to that of vector control and WT FLOT2 (Fig. 6*D*). Although prior experiments had used HeLa as a model, we used HEK293T as these cells are more suitable for transient overexpression of multiple proteins.

Overexpression of WT FLOT2 resulted in a reduction in EGFR phosphorylation (normalized to EGFR levels) upon EGF stimulation (Fig. 6*E*). The same effect was observed when the Y163F mutant (which associates with the plasma membrane) was overexpressed. However, when the G2A mutant (unable to associate with the plasma membrane) was expressed, EGF-induced EGFR phosphorylation (normalized to EGFR levels) was comparable to that of vector control. Moreover, only weak co-immunoprecipitation of the G2A mutant was observed when the EGFR was immunoprecipitated compared to either WT or Y163F FLOT2 (Fig. 6*D;* compare lanes 7-9 to lanes 4-6 or 10-12). These findings indicated that FLOT2 requires membrane association by glycine myristoylation for its effects on EGFR signaling.

### FLOT2 overexpression increases EGFR levels and reduces EGFR signaling in PC9 cells

We generated stable clones expressing empty vector (EV) or WT FLOT2 in PC9 cells, which endogenously express significantly less FLOT2 than HeLa or H441 cells (Fig *S4C*). Considering these cells have less endogenous FLOT2, it is easier to interpret the effects of FLOT2 overexpression. We observed that overexpression of WT FLOT2 increased total EGFR levels, and also reduced both EGFR and MAPK activation (Fig. 6*F*). Two separate WT clones were compared to the average of two EV clones for analysis.

### FLOT2 inhibits EGFR-dependent anchorage dependent and independent cell growth *in vitro*

Next, we tested whether the modulation of EGFR activity by FLOT2 contributed to changes in cancer cell growth. Growth of FLOT2 KO HeLa cells was compared to that of control cells by counting viable cells over a span of a week with acridine orange and propidium iodide (AOPI) staining. There was a statistically significant increase in growth in the FLOT2 KO HeLa cells (Fig. 7*A*). We next measured growth with EGF +/- erlotinib in both the control and the FLOT2 KO HeLa cells. 1 µ M erlotinib was used since this concentration did not affect unstimulated HeLa growth, however it was found to completely inhibit EGF-stimulated growth (Fig. S9*A* and S9*B*). The increase in growth in FLOT2 KO in the absence of exogenous EGF is partially inhibited by erlotinib, while the EGF stimulated increase in growth is completely inhibited in erlotinib (Fig. 7*B* and S9*B*). This is consistent with the increased growth in FLOT2 KO cells being driven partially by EGFR activity. Similar to HeLa, silencing of FLOT2 in H441 cells resulted in increased growth compared to control H441 cells (Fig. 7*C*). The growth of the EGF stimulated control H441 cells was inhibited by erlotinib but the growth of the control H441 cells in the absence of EGF was not inhibited (Fig. 7*D*). Both the increased growth of unstimulated and EGF stimulated FLOT2 KD cells was completely inhibited by erlotinib (Fig. 7*D*). This is consistent with the increased growth of H441 FLOT2 KD cells with or without EGF being driven predominantly by the activated EGFR. Together, the data from these two models indicates that the increased growth of the H441 and HeLa cells lacking FLOT2 is due, at least in part to, an increase in EGFR activation. To confirm that the growth effects were not due to off-target or non-specific siRNA effects, three different siRNA’s for FLOT2 increased growth in both HeLa (Fig. S9*C*) and H441 (Fig. S9*D*). Noteworthy, both HeLa and H441 express multiple EGFR ligands at the mRNA level which predicts autocrine signaling in both cell lines (Table S2).

**Figure 7.**
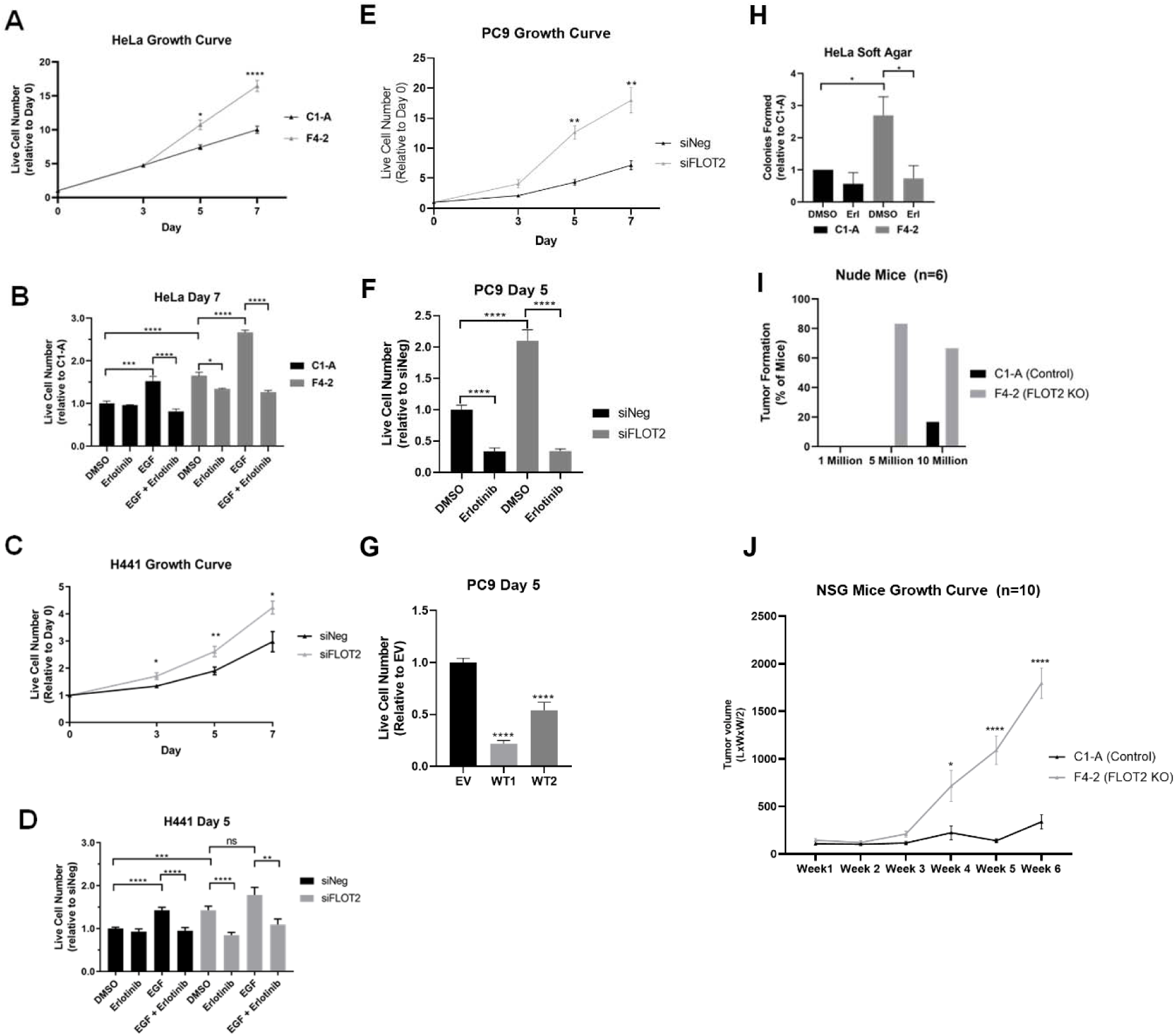
FLOT2 knockout/knockdown increases EGFR dependent *in vitro* growth, anchorage-independent growth, and tumor formation and growth in mice, while FLOT2 overexpression decreases *in vitro* growth. (A) Control (C1-A) and FLOT2 KO CRISPR (F4-2) HeLa cells were plated and counted with AOPI staining for viable cells on Day 0, 3, 5 and 7. Cell number was normalized to the Day 0 counts of each cell line, and the graph shows the average (± SEM) of three independent experiments. Asterisks (*) denotes p <0.05 and (****) denotes p <0.0001 when comparing the corresponding time points for C1-A and F4-2 using Student’s *t*-test. (B) Control (C1-A) and FLOT2 KO CRISPR (F4-2) HeLa cells were plated and treated with DMSO (control), 1 µ M erlotinib, 25 ng/mL EGF, or erlotinib + EGF, and subsequently counted with AOPI staining for viable cells on Day 7. Cell number was normalized to the Day 0 counts of each cell line, and then normalized relative to C1-A treated with DMSO. The graph shows the average (± SEM) of three independent experiments. Asterisks (*) denotes p <0.05, (***) denotes p <0.001 and (****) denotes p <0.0001 using Student’s *t*-test. (C) FLOT2 was knocked down in H441 cells by specific siRNA for 48 hours, and then re-plated for cell counts. Cells were counted with AOPI staining for viable cells on Day 0 (day following re-plating), 3, 5 and 7. Cell number was normalized to the Day 0 counts of each transfection, and the graph shows the average (± SEM) of three independent experiments. Asterisks (*) denotes p <0.05 and (**) denotes p <0.01 when comparing the corresponding time points for non-targeting siRNA (siNeg) and siFLOT2 using Student’s *t*-test. (D) FLOT2 was knocked down in H441 cells by specific siRNA for 48 hours, and then re-plated for cell counts. The day following re-plating, cells were treated with DMSO (control), 1 µ M erlotinib, 25 ng/mL EGF, or erlotinib + EGF, and subsequently counted with AOPI staining for viable cells on Day 5. Cell number was normalized to the Day 0 counts of each transfection, and then normalized relative to siNeg. The graph shows the average (± SEM) of five independent experiments. Asterisks (**) denotes p <0.01, (***) denotes p <0.001, (****) denotes p <0.0001, and non-significant (ns; p >0.05) using Student’s *t*-test. (E) FLOT2 was knocked down in PC9 cells by specific siRNA for 48 hours, and then re-plated for cell counts. Cells were counted with AOPI staining for viable cells on Day 0 (day following re-plating), 3, 5 and 7. Cell number was normalized to the Day 0 counts of each transfection, and the graph shows the average (± SEM) of three independent experiments. Asterisks (**) denotes p <0.01 when comparing the corresponding time points for non-targeting siRNA (siNeg) and siFLOT2 using Student’s *t*-test. (F) FLOT2 was knocked down in PC9 cells by specific siRNA for 48 hours, and then re-plated for cell counts. The day following re-plating, cells were treated with DMSO (control) or 100 nM erlotinib and subsequently counted with AOPI staining for viable cells on Day 5. Cell number was normalized to the Day 0 counts of each transfection, and then normalized relative to siNeg. The graph shows the average (± SEM) of four independent experiments. Asterisks (****) denotes p <0.0001 using Student’s *t*-test. (G) Stable PC9 cells expressing empty vector (EV) or overexpressing wild-type FLOT2 (WT) were plated and counted with AOPI staining for viable cells on Day 0 and Day 5. Cell number was normalized to the Day 0 counts of the average EV counts, and the graph shows the average (± SEM) of three independent experiments. Asterisks (****) denotes p <0.0001 using Student’s *t*-test. (H) Control (C1-A) and FLOT2 KO CRISPR (F4-2) HeLa cells were suspended in soft agar and plated. The top layer was fed with media containing DMSO (control) or 1 µ M erlotinib for three weeks. Following treatment, colonies formed were counted and plotted relative to C1-A DMSO, with the average (± SEM) of three independent experiments. Asterisks (*) denotes p <0.05 following Student’s *t*-test. (I) Control (C1-A) or FLOT2 KO CRISPR (F4-2) HeLa cells were orthotopically injected into the flank of 6 nude mice/group at concentrations of 1, 5 and 10 million cells per flank. Tumors were measured for 10 weeks, and tumor formation was considered as a tumor over 4mm^3^. The percentage of mice from each condition which formed tumors were plotted. (J) Control (C1-A) or FLOT2 KO CRISPR (F4-2) HeLa cells were suspended in a 1:1 ratio of media:Matrigel at 5 million cells per mouse, and injected orthotopically into the flank of 10 NSG mice per cell line. Tumor volume was tracked over 6 weeks, and plotted as average tumor volume. Asterisks (*) denotes p <0.05, (****) and denotes p <0.0001 when comparing C1-A to F4-2 average tumor volume using Student’s *t*-test.

To further confirm the effects of FLOT2 on cancer cell growth, we determined that FLOT2 KD in PC9 cells significantly increases the cell lines growth over time. This effect was EGFR-dependent, as 100 nM of erlotinib inhibited the growth of the siNEG treated control cells and reduced the increase in growth seen in FLOT2 KD cells (Fig. 7*E* and *F*). Overexpression of WT FLOT2 in PC9 cells significantly reduced cell growth, providing evidence that FLOT2 negatively regulates growth in these EGFR-driven cells (Fig. 7*G*).

We next examined if FLOT2 also regulates anchorage-independent growth, which is considered a major property of transformed cells. Over a span of three weeks, we observed increased anchorage-independent colony formation in FLOT2 KO HeLa which was completely reversed by erlotinib treatment (Fig. 7*H*). Colonies were counted as >100 µ m in size under a microscope. While the growth of FLOT2 KO cells in 2D culture was partially inhibited by erlotinib, the growth in soft agar was completely inhibited, suggesting a more important role for EGFR in anchorage independent growth. In summary, FLOT2 negatively regulates EGFR-dependent anchorage dependent and independent cell growth *in vitro*.

### FLOT2 negatively regulates tumor formation and growth in nude and NSG mice, respectively

We next tested if FLOT2 negatively regulates EGFR-dependent tumor growth *in vivo.* In a pilot study we injected varying numbers of control or FLOT2 KO HeLa cells into the flank of nude mice and conducted weekly measurements to track tumor formation. There was no tumor formation with 1 million cells injected in either group, however both the 5 and 10 million cell groups exhibited increased fraction of mice with tumors in the FLOT2 KO HeLa compared to control (Fig. 7*I*). 5 out of 6 mice in the FLOT2 KO 5 million cell group, and 4 out of 6 mice in the FLOT2 KO 10 million cell group developed tumors. In contrast, no mice in the 5 million cell control group, and only one mouse in the 10 million cell control group, developed tumors. Following injection in nude mice there was an initial decrease in the injected tumor volume, followed by a four-week latency period, and then slow tumor growth. We speculated that there may have been an innate immune response leading to initial tumor loss and dormancy, therefore we next tested growth in NOD SCID gamma (NSG) mice. Five million cells were injected in NSG mice and tumor volume was tracked over six weeks. There was a statistically significant increase in tumor volume in the FLOT2 KO group compared with control (Fig. 7*J*). Thus, loss of FLOT2 results in increased tumor formation and growth *in vivo*.

## Discussion

We determined that FLOT2 negatively regulates EGFR activation, dimerization, trafficking, signaling, and cancer cell growth in HeLa, H441 and PC9 cancer cell lines. Based on our data, we hypothesize that EGFR is activated by increased receptor dimerization (Fig. 5 *A-D*) upon the depletion of FLOT2, which in turn leads to enhanced signaling through the EGFR/MAPK pathway, resulting in increased cell proliferation and tumor growth (Figs. 1 *C* and *D*, 2*A, B* and *D*, 3*A, B*, and *C*, 7 *A-J*, and S4 *A* and *B*, and S9*B-D*). We also found that elevated activation of EGFR in the absence of FLOT2 leads to increased Cbl recruitment (Fig. 2*C*), EGFR ubiquitination (Figs. 2*C* and 3*D*), and EGFR endocytosis and Cbl-mediated degradation (Figs. 1*E,* 4 *A-D* and 5 *E-G*). This downregulation of EGFR and dimerization is not occurring via lipid-rafts, as cholesterol depletion did not prevent dimerization or downregulation (Fig S6*E-F*), and the downregulation of EGFR relies on the lysosomal pathway (Fig. 4*E*). Consistent with these observations, when the EGFR inhibitor erlotinib was used, the constitutive increase in phosphorylation of EGFR and MAPK seen upon FLOT2 depletion was downregulated (Fig. 2*B,* 3*C).* However, cabozantinib and defactinib, inhibitors of other RTK’s including MET, VEGFR, RET, KIT, AXL, and FAK, did not affect MAPK phosphorylation (Fig. S5). Further, erlotinib reduced the anchorage dependent and independent growth seen upon FLOT2 depletion (Fig. *7B, D, F* and *H*).

In the present study, we developed a novel approach which included a primary SILAC MS screen to identify proteins that interacted with Cbl with and without EGFR activation, followed by an inexpensive, fast and sensitive secondary siRNA screen coupled to in-cell pEGFR ELISA which was specifically designed to identify proteins that affect EGFR activation/degradation (Fig. 1*A* and *B*, S1*A* and *C*, Table 2, Table S1). The most significant proteins identified by SILAC had no effect on EGFR phosphorylation in the ELISA screen. Of note, a similar study determining the EGF-induced interactome of Cbl and Cbl-b was recently published which used proteomics to study complex formation, however this study failed to observe flotillin interaction with Cbl (27).

We discovered that for some proteins identified in our SILAC screen, the changes upon EGF stimulation could not be generated due to the complete absence of the corresponding peptides in one of the groups. These data, however, might be very useful for future studies as it could indicate an all-or-none response to treatment. Surprisingly, EGFR itself was not found in the complex with Cbl in our SILAC screen. This could be due to poor trypsin digestion or insufficient ionization of EGFR peptides in the mass spectrometer. Alternatively, this might indicate the transient nature of EGFR/Cbl interaction. Consistent with this speculation, a recent EGFR MS screen failed to identify Cbl (28).

Not all published work confirms our observations. A study in hepatocellular carcinoma discovered that FLOT2 overexpression increased cell proliferation and migration, as well as ERK activation, and another study in breast cancer determined FLOT2 KD reduced cell growth (29, 30). In intrahepatic cholangiocarcinoma, FLOT2 overexpression was correlated with lymph node metastasis, and in nasopharyngeal carcinoma FLOT2 overexpression increased proliferation, and expression was associated with metastasis (31–33). Further, FLOT1 and/or FLOT2 are overexpressed in many cancers including breast, esophageal squamous cell, hepatocellular, gastric, lung, nasopharyngeal, oral squamous cell carcinomas and melanoma; their overexpression is associated with poor prognosis and reduced patient survival in some of these cancers (34). These studies, along with the fact that FLOT1 or FLOT2 KO in mice have no obvious phenotype, have led to suggestions that FLOT1 or FLOT2 inhibition could be an effective treatment strategy (10).

Although some prior studies have seen FLOT2 overexpression correlating with poor outcomes, we suggest that the data and interpretation is more nuanced. We performed transcriptome analysis using The Cancer Genome Atlas Program (TCGA) and found that high FLOT2 expression had a statistically significant better disease free survival in invasive breast carcinoma and kidney renal papillary cell carcinoma (Fig. S10A), but had statistically significant worse disease free survival in brain lower grade glioma, adrenocortical carcinoma, and stomach adenocarcinoma (Fig. S10B). The rest of the data sets in TCGA, including lung squamous cell carcinoma, lung adenocarcinoma, cervical squamous cell carcinoma and endocervical adenocarcinoma (Fig S10C), did not show statistically significant differences in outcome. Overall, this supports our model that FLOT2 expression alone does not predict the effects of FLOT2 on cancer outcomes and is not enough to support molecular targeting of FLOT2 for drug therapy. The role of FLOT2 will require studies of the biochemical effects of FLOT2 on cell signaling in different cancer types.

It is important to note, however, many of these studies did not explore the role of EGFR in changes in growth, migration, or prognosis as it relates to FLOT2 protein function, as they only look at gene expression. Our model indicates that FLOT2 inhibits EGFR activation, resulting in growth inhibition. However, it is plausible that this is occurring in the aforementioned studies that investigated FLOT2, but due to the difference in cell types experimented upon, there are alternative pathways being affected which override the effects on EGFR. These studies as well as ours underscore the hypothesis that FLOT2 may regulate oncogenic signaling differently depending on context. There is significant heterogeneity between different cancer cell types, and it is likely that various mutations or altered signaling pathways could affect the role of FLOT2. Thus, FLOT2 inhibition may only be successful in certain cell types where FLOT2 correlates with poor survival, such as brain lower grade glioma, adrenocortical carcinoma, and stomach adenocarcinoma, whereas inhibition in cancers such as breast invasive carcinoma and kidney renal papillary cell carcinoma, where FLOT2 expression seems to improve survival, may be detrimental (Fig S10 *A* and *B*). Our work indicates that further experimentation is required to fully characterize the range of mechanisms that FLOT2 affects and why certain cell types respond differently to FLOT2 modulation.

Although some studies contradict our data, others support it. We found that FLOT2 depletion led to increased activation of EGFR in the presence and absence of exogenous ligand, increased downstream signaling, accelerated *in vitro* cell growth, increased anchorage independent growth and increased tumor formation *in vivo* (Figs. 2 *A-C*, 3 *A-D* and 7 *A-J*). Several studies have reported similar findings, including an increase in HeLa cell proliferation and MAPK activation (35), increased EGFR phosphorylation and MAPK activation in MCF7 (19), and EGFR-dependent adherens junction formation and motility changes in A431 cells with FLOT2 KD (18).

We speculate that the reported association between FLOT2 overexpression and poor patient survival might indicate that FLOT2 is merely a marker, not the driver. A majority of studies investigating the oncogenic role of FLOT2 are studying correlation to gene expression, and not mechanistic data. For example, the *FLOT2* gene, located on chromosome 17 within the *ERBB2* amplicon, is amplified in HER2 amplified cancers and a positive correlation exists between FLOT2 and HER2 expression in breast (36) and gastric cancer tissues (37). We propose that the field moving forward focuses on more mechanistic insight, instead of expression profiles, into the role of FLOT2 in cancer proliferation and migration, and determination of why different models respond differently to FLOT2 modulation.

The decreased steady-state levels of EGFR observed with siRNA mediated KD and CRISPR mediated KO of FLOT2 are partially explained by enhanced Cbl-mediated protein turnover due to increased constitutive activation of EGFR. Phosphorylated EGFR is recognized by Cbl ubiquitin ligases and is quickly targeted for degradation to prevent prolonged signaling and uncontrolled cell proliferation (38). The association between Cbl and EGFR is increased when FLOT2 is lost (Fig. 2*C*), which could be explained in a variety of ways. Loss of FLOT2 could result in increased EGFR activation which would subsequently lead to increased association of Cbl to activated EGF. Alternatively, FLOT2 interactions with EGFR and/or Cbl could shield EGFR from Cbl, and loss of FLOT2 would result in greater structural access for Cbl to EGFR. Future work is required to fully understand how FLOT2 prevents Cbl recruitment to the activated EGFR. Regarding EGFR trafficking following ubiquitination by Cbl, a recent study concluded that flotillins in lipid raft microdomains play a role in driving EGFR into multivesicular endosomes, forming intraluminal vesicles, and resulting in transport that is independent of endosomal sorting (39), although in our study lipid raft disruption with simvastatin did not reduce EGFR dimerization (Fig. S6*F*). Although our study showed endosomal trafficking by EEA1 co-localization (Fig. 5 *E* and *F*), it is possible that FLOT2 plays a role in intraluminal vesical transport as well, and future studies should consider this possibility.

The restoration of the EGFR protein levels in FLOT2 KD/KO cells seen upon KD of Cbl and Cbl-b further confirm that EGFR down-regulation is Cbl-dependent (Fig. 4 *A-D*). In agreement with our data, a recent study has reported on the significant positive association between FLOT2 and EGFR protein expressions in non-small cell lung cancer, although the study postulated that high FLOT2 expression correlates with poor overall survival, which contradicts our data (40). Noteworthy, siRNA mediated depletion of FLOT2 decreased steady-state levels of ErbB2 and ErbB3 in MCF7 and SKBR3 cells (20, 36). Given the homology among ErbB family members, it would be important to test if this decrease of steady state protein levels was also accompanied by the increase in their phosphorylation and down regulation. To our knowledge, the effect of FLOT2 on EGFR ubiquitination has not been previously reported in the literature.

Our data show a constitutive interaction between EGFR and FLOT2 based on co-immunoprecipitation. How FLOT2 physically interacts with EGFR remains unanswered. Using the FLOT2 G2A mutant, which cannot be myristoylated and recruited to the membrane, we demonstrated that FLOT2 association with the membrane was necessary for both interaction with and inhibition of the EGFR (Fig. 6, *D* and *E*). However, it is unclear whether membrane association without interaction with EGFR would be sufficient to reduce EGFR phosphorylation. Although we discovered a statistically significant increase in EGFR dimerization upon loss of FLOT2 in both HeLa and H441 cells (Fig. 5 *A-D*), it is unclear how FLOT2 regulates EGFR dimerization and activation. One possibility is that by binding to EGFR, FLOT2 directly interferes with dimerization of the EGFR or that FLOT2 prevents EGFR from being in an “activatable” state, and loss of FLOT2 allows endogenous ligands to activate and dimerize the receptor. For example, it is known that H441 cells express both EGF and tumor growth factor (TGF) alpha, and FLOT2 may allow EGFR to be activated by these ligands (41). We also show that both HeLa and H441 express multiple EGFR ligands (Table S2). This is consistent with our data showing that HeLa and H441 growth increases upon FLOT2 loss and that growth is blocked by EGFR inhibition (Fig 7*B* and *D)*.

Another possibility is that FLOT2 affects lipid rafts. When we inhibited cholesterol synthesis with simvastatin, which should disrupt lipid raft composition, we did not observe any rescue on EGFR downregulation by FLOT2 (Fig. S6*E*). Further, lipid raft disruption by simvastatin treatment actually induced further dimerization in FLOT2 KO HeLa cells (Fig. S6*F*). Interestingly, a previous study indicated that cholesterol depletion enhanced EGFR clustering at the cell membrane, which supports our finding that cholesterol depletion increased EGFR dimerization (42). Considering that FLOT2 has been used as lipid raft markers (10, 15), it is possible that loss of FLOT2 also depletes lipid raft formation, and may cause EGFR clustering and dimerization in a similar manner to cholesterol depletion. Future experiments may help better define this role.

Since we observed increased EGFR activation even in the absence of exogenous EGF, it is possible that FLOT2 reduces the production/shedding of endogenous EGFR ligands. In agreement with this possibility, depletion of FLOT2 was found to increase mRNA levels of HB-EGF in the lungs of FLOT2 KO mice (35). Alternatively, FLOT2 may inhibit the activity of a disintegrin and metalloprotease (ADAM) family of proteins, responsible for the shedding of the EGFR ligands, such as amphiregulin or tumor growth factor tumor growth factor TGFα (43). This inhibitory action has been so far only demonstrated for FLOT1 on ADAM10 activity in *Xenopus* model of neuronal development (44). This is all speculative, and certainly warrants future investigations into the mechanistic effects of FLOT2 on EGFR activation.

In conclusion, our novel approach to study EGF-induced complexes on Cbl identified FLOT2 as a negative regulator of EGFR activation. Our findings combined with previous reports provide evidence that FLOT2 may have varying roles in carcinogenesis depending on cell type and justifies the need for further research to clarify the mechanisms by which FLOT2 regulates EGFR (and other RTK) activation and cancer cell growth.

## Materials and Methods

### Materials

Dulbecco’s modified Eagle’s medium (DMEM), RPMI and fetal bovine serum (FBS) were obtained from Invitrogen (Carlsbad, CA). Dulbecco’s phosphate buffered saline (DPBS) was purchased from Mediatech Inc. (Herndon, VA). LB broth (BLF-7030) was from KD Medical (Columbia, MD). Sodium orthovanadate was from Fisher Chemicals (Fairlawn, NJ). NP40 Cell Lysis Buffer (FNN0021), Geneticin Selective Antibiotic (G418 Sulfate, 10131027) and OptiMEM medium (31985062) were from ThermoFisher Scientific (Waltham, MA). Puromycin dihydrochloride (P9620) and leupeptin (L9783) were from Sigma Aldrich (St. Louis, MO). Recombinant human EGF was purchased from BD Biosciences, Inc. (San Jose, CA). Erlotinib (S1023), bortezomib (S1013), MG132 (S2619), chloroquine (S6999), cabozantinib (S1119), defactinib (S7654) and simvastatin (S1792) was from Selleckchem (Houston, TX). Tissue culture plastic ware and other laboratory consumables were purchased from commercial sources.

### Antibodies

Anti-ubiquitin (P4D1, sc-8017), anti-Cbl (C-15, sc-170), anti-Cbl-b (G-1, sc-8006), anti-ERK2 (D-2, sc-1647) and anti-HSC70 (sc-7298) antibodies were purchased from Santa Cruz Biotechnology (Dallas, TX). Anti-pMAPK T202/Y204 (9101), anti-Flotillin-2 (C42A3, 3436), anti-Flotillin-1 (3253), anti-pMET Y1234/1235 (3077), anti-MET (3127), anti-LC3A/B (12741) and anti-myc-tag (2278)antibodies were from Cell Signaling Technology (Danvers, MA) and used for immunoblotting. Phospho-EGFR pY845 (44-784G), Anti-pFAK Y397 (44624G) and anti-EGFR pY1173 (44-794G) were from ThermoFisher Scientific (Waltham, MA). Anti-FLAG (A8592) and anti-FAK (06-543) antibodies were obtained from Sigma Aldrich. Anti-phosphotyrosine (4G10, 05-321) antibodies were obtained from Millipore (Billerica, MA). Anti-calreticulin antibody (ab2907) were obtained from Abcam (Cambridge, UK). Anti-EGFR antibodies (199.12, MA5-13319) were purchased from ThermoFisher Scientific and used for immunoprecipitation. Anti-EGFR (2232) and anti-phospho-EGFR pY1068 (2234) antibodies were obtained from Cell Signaling Technology and used for immunoblotting. anti-FLOT2 (B-6) antibody was obtained from Santa Cruz biotechnology and used for immunoprecipitation. Anti-EGFR (528, sc-120) from Santa Cruz Biotechnology and EEA1 (3288) and LAMP1 (D2D11) from Cell Signaling Technology were used as primary antibodies for immunofluorescence, and secondary antibodies Alexa Fluor 488 (715-545-150) and 594 (111-585-144) were purchased from Jackson ImmunoResearch (West Grove, PA). Anti-EGFR (H11, MA5-13070) was purchased from ThermoFisher Scientific and used for proximity ligase assays.

### Cell lines

The human embryonic kidney (HEK293T) and HeLa (CCL2) cells were obtained from ATCC (Manassas, VA) and maintained in culture in DMEM supplemented with 10% FBS. H441 cells were obtained from ATCC and maintained in culture in RPMI medium with 10% FBS. PC9 cells were obtained from Sigma-Aldrich maintained in culture in RPMI medium with 10% FBS. All cell lines were grown at 37°C, 5% CO_2_ and high humidity. All cell lines including stable clones were routinely evaluated for mycoplasma contamination using the LookOut Mycoplasma PCR Detection Kit (MP0035, Sigma-Aldrich) according to the manufacturer’s protocol.

### siRNA transfections

siRNA transfections were performed as previously described (24). When indicated, transfected H441 cells were treated with 10 M erlotinib for 24 hours before harvesting. siRNAs for FLOT1 (s19913, s19914, s19915), FLOT2 (s5284, s5285, s5286), Cbl (s2476), Cbl-b (s2479) and negative control (4390844) all were purchased from ThermoFisher Scientific.

### Immunoblotting and immunoprecipitation

Proteins were harvested as previously described (24). For immunoprecipitation, transfected HeLa, H441 or HEK293T lysates containing 1 mg protein were incubated with anti-EGFR (199.12) or anti-Cbl (C-15), or anti-FLOT2 (B-6) antibody and Protein A/G PLUS agarose beads (sc-2003) all from Santa Cruz Biotechnology overnight at 4°C with tumbling. Immunoblotting was performed as previously described (24). m-IgG_κ_ BP-HRP secondary antibody (sc-516102, Santa Cruz Biotechnology) was used for immunoblotting detection for anti-FLOT2 IP. Each experiment was repeated at least 3 times. Densitometric analysis of immunoblot band intensities was performed using Adobe Photoshop software version CC 2017 (Adobe Systems Inc., U.S.A). Data were presented as an average ± SEM.

### SILAC mass spectrometry

HeLa cells were cultured in DMEM SILAC lysine and arginine-free medium (#88364, ThermoFisher Scientific) supplemented with 10% FBS. Two-state SILAC media were prepared to contain different stable-isotope labeled versions of arginine (0.398 mM) and lysine (0.798 mM). All heavy isotope labeled amino acids (99% pure) were purchased from Cambridge Isotope Laboratories (Andover, MA). Media were prepared containing non-labeled arginine and lysine (light), and ^13^C_6_^15^N_4_ arginine (#CNLM-539) and ^13^C_6_^15^N_2_ lysine (#CNLM-291) (heavy). The labeling efficiency was evaluated with MS after five passages of HeLa cells in the labeling media. Upon achieving >97% labeling efficiency, Hela cells were plated on 150 mm dishes to reach 90% confluency, starved for 5 hours and treated as follows: no EGF stimulation (light), and stimulation with 100 ng/mL EGF for 30 min (heavy). Upon treatments, the cells were lysed in Lysis buffer (10 mM TrisHCl, 150 mM NaCl, 5 mM EDTA, 1% Triton X-100, 10% Glycerol) containing 1 mM Sodium Vanadate and the protease inhibitor cocktail Complete Mini (Roche). Then, Cbl was immunoprecipitated from 20-50 mg of total protein from each treatment using anti-Cbl AB coupled to agarose beads (sc-170 AC, Santa Cruz Biotechnology). The immunoprecipitated proteins were eluted from Cbl AB-agarose beads with Pierce IgG Elution Buffer (21004, pH 2.8) and the pH was adjusted to 7 with 1M Tris buffer (pH 8). The eluate was then transferred to the Amicon Ultra-4 centrifugal filter units 10 kDa (UFC801024, Millipore) to reduce the volume and to exchange the buffer for 20 mM HEPES. The immunoprecipitated proteins from three treatments were mixed and either denatured by 9 M Urea/20 mM HEPES buffer or mixed with 2x Laemmli Sample Buffer (1610737, Bio-Rad) and separated by SDS-PAGE to analyze individual fractions. In both cases the samples were reduced with DTT, alkylated with iodoacetamide and digested with modified sequencing grade Trypsin (Promega, Madison, WI) at 30°C for 16 h. Mass spectrometry analysis of tryptic peptides was performed on an LTQ-Orbitrap Elite (Thermo Scientific, San Jose, CA) mass spectrometer interfaced with an Easy-nLC 1000 (Thermo Scientific, San Jose, CA) as previously described (45). Data analysis was performed using the MaxQuant software package (version 1.4.0.3) with the Andromeda search engine (46). MS/MS spectra were searched against the UniProt human protein database (May 2013, 38523 entries) and quantification was performed using default parameters for three state-SILAC in MaxQuant as previously described (45).

### In-cell phospho-EGFR ELISA screen

Pooled siRNAs for each gene were mixed with Lipofectamine RNAiMAX Transfection Reagent (13778150, ThermoFisher Scientific) in Opti-MEM medium and added to the trypsinized HeLa cells. HeLa cells were then plated on 96-well plates at 10,000 cells/well and incubated for 24 hours. After the initial incubation, the medium was changed, and the cells were incubated for an additional 24 hours. Each gene was knocked down by the mixture of 3 siRNA oligos at 20 nM (Table 2) and analyzed in triplicates. HeLa cells transfected with siRNA targeting Cbl and Cbl-b at 10 nM each were used as a positive control for each 96-well plate analyzed (Fig. S1, *A* and *B*). Non-targeting siRNA was used as a negative control (siNeg). Before the analysis, transfected HeLa cells were starved for 2 hours in FBS-free DMEM medium and then stimulated with 100 ng/mL EGF for up to 30 minutes. Upon treatment the cells were fixed in 4% formaldehyde and in-cell ELISA was performed according to the manufacturer’s protocol (kit 62205, ThermoFisher Scientific). pEGFR antibodies provided in the kit were replaced with pEGFR Y1173 antibodies (44-794G) from ThermoFisher Scientific. Total EGFR levels were not measured. Janus Green Whole-Cell staining was performed to assess the total cell number in each well. The results were calculated as a ratio of absorbance values from pEGFR staining normalized to the absorbance values from the Whole-Cell staining.

### Plasmid transfection to HEK293T cells

Calcium phosphate (ProFection E1200, Promega Corp., Madison, WI) was used for plasmid transfection into HEK293T cells. Eighteen hours post transfection, medium on transfected cells was changed. At 48 hours post transfection, cells were starved without serum for 3 hours and then stimulated with 100 ng/mL EGF for up to 15 minutes, harvested and lysed. Each experiment was repeated at least 3 times. Plasmids for overexpression in HEK293T cells were purchased from OriGene (Rockville, MD): pCMV6 (PS10001), FLOT1 (RC200231), FLOT2 (RC220884). The use of HA-tagged Cbl plasmid in pCEFL expression vector as well as WT EGFR in pcDNA3 vector has been previously reported (47). Site-directed mutagenesis using QuikChange II Site-Directed Mutagenesis Kit (Stratagene, La Jolla, CA) was performed to create the FLOT2 G2A and FLOT2 Y163F mutants. All the constructs were confirmed by DNA sequencing.

### CRISPR knockout of FLOT2 in HeLa cells

Flotillin-2 protein was knocked out using a CRISPR/cas9 system to disrupt the *FLOT2* gene as previously described (48). A guide sequence was designed to target the fourth exon of the gene by annealing the following two oligos (5’ – CACCGAAACGTCGTCCTGCAGACCC – 3’ & 5’ – AAACGGGTCTGCAGGACGACGTTTC – 3’) and cloning them into the sgRNA scaffold of a CRISPR/cas9 vector using BbsI restriction sites [pSpCas9(BB)-2A-Puro (PX459) V2.0 was a gift from Feng Zhang (Addgene plasmid #62988)]. Control sequences were similarly constructed to target eGFP (5’ – CACCGGCACTACCAGAGCTAACTCA – 3’ & 5’ – AAACTGAGTTAGCTCTGGTAGTGCC – 3’). The resultant plasmids were transformed into Stbl3 chemically competent E. coli, after which they were purified using a Qiagen Plasmid Maxi Kit. Following verification of the purified plasmids’ sequences, 1 g was transfected into HeLa cells, and the cells were incubated at 37°C/5% CO_2_ for 2 days. The medium was replaced with fresh growth medium for 24 hours to allow the cells to recover. The growth medium was then replaced with drug-selective medium (2 g/mL puromycin) for one week to select for cells that had taken up the CRISPR/cas9 plasmid. Following selection, the remaining cells were counted and plated in 96-well plates for limiting dilutions. Individual clones derived from this targeting were then expanded and tested for the absence of FLOT2 protein as well as the presence of genetic defects at the *FLOT2* locus. To assess the unique genetic modifications of FLOT2 KO clones, DNA from the cells was extracted according to the protocol for DNeasy Blood & Tissue Kit (69504, Qiagen). Following extraction, template DNA was amplified using primers directed to the exon 4 locus (F: 5’ – agaggcttcagacagattccag – 3’, R: 5’ – cccttcactcataccctctcc – 3’). PCR products were run on a gel to confirm the presence of a 453 bp band and subsequently run through a PCR cleanup protocol (QIAquick PCR Purification Kit, 28104). The above forward primer was used for sequencing. Chromatograms were visualized using the SnapGene Viewer program and compared to parental HeLa cells to determine genetic changes to *FLOT2*. Clone F4-2 was found to have a homozygous nine-base-pair deletion, while clone F4-20 showed a distorted nucleotide sequence after the DNA cut site (Fig. S3*A*). Both F4-2 and F4-20 demonstrated the complete loss of FLOT2 protein while clones derived from the control sequences had wild-type sequences of *FLOT2* and full-length protein.

In the F4-2 clone, a homozygous deletion of 9 base pairs (ACC CTG GAG) from Exon 4 (predicted to result in the loss of the 3 amino acids TLE) in positions 106-108 led to the complete KO of FLOT2 protein (Fig. 3*A*). We speculate that the deletion of 9 base pairs might have led to changes in FLOT2 mRNA, leading to its instability (49). In the F4-20 clone, the introduced double-strand break resulted in the complete disruption of the wild-type sequence of the *FLOT2* gene (Fig. S3*A*) and loss of the protein (Fig. 3*A*).

### Generation of HeLa and PC9 stable clones

HeLa cells were transfected with 20 g of empty vector pCMV6 (PS10001) or 18 g of FLOT2 (RC220884) plasmid with Lipofectamine LTX Plus Reagent (15338100, ThermoFisher Scientific) according to the manufacturer’s instructions. The medium was changed 24 hours post transfection to allow the cells to recover. After 24 hours, the cells were split into selection medium containing 0.4 mg/mL Geneticin (G418) and allowed to grow until visible colonies were formed, changing media every 3-4 days as needed. Several clones were analyzed for the expression of FLAG-FLOT2. The control clones were selected based on their resistance to G418. The positive clones were routinely maintained in 0.1 mg/mL of G418 to prevent loss of the transgene.

PC9 cells were transfected in a 6-well plate with 1 µ g empty vector pCMV6 or FLOT2 plasmid for 24 hours, and then media was changed, according to the manufacturer protocol for Lipofectamine 3000 (ThermoFisher L3000001). 2.5 mg/mL G418 was then used for selection until visible colonies formed from single cell clones. Clones were expanded and analyzed for myc-tag.

### Immunofluorescence and confocal microscopy

Cells were plated and stained with EGFR (1:50) and EEA1 (1:100) as previously described (50). Briefly, cells were seeded on coverslips, serum starved for 4 hours, then stimulated with 25 ng/mL EGF for 15 minutes and then fixed with 2% formaldehyde. They were then washed, blocked with PBS/FBS, permeabilized with 0.1 % saponin and then treated with primary followed by secondary antibody. Coverslips were washed and mounted on slides with Fluoromount-G (0100-01) from SouthernBiotech (Birmingham, AL)

Images were acquired using Leica SP8 inverted confocal laser scanning microscope using 63x/1.4 objective. Co-localization analysis was performed as previously described (51). Briefly, images were manually thresholded with ImageJ and the multiply feature in the image calculator generated an image with overlapping pixels (EGFR and EEA1 or EGFR and LAMP1), and the overlapping intensity was divided by the intensity of EGFR to calculate percent co-localization. Experiments were performed three times, and representative images are shown.

### Proximity ligation assay

EGFR mAb was conjugated separately to Duolink In Situ Probemaker PLUS (DUO92009) and Duolink In Situ Probemaker MINUS (DUO92010) as described by the manufacturer instructions (Sigma Aldrich). Dimerization of PLUS and MINUS probes was performed according to the manufacturers protocol. Briefly, cells were fixed in 2% formaldehyde, permeabilized with 0.1% Triton for ten minutes each, and blocked with PLA Blocking Solution for thirty minutes. EGFR/probe conjugates were diluted 1:25 each into PLA probe diluent, and added to cells plated on cover slips, encircled by hydrophobic pen ink to keep the reaction on the cover slip. Antibodies were incubated overnight at 4 degrees Celsius, then washed with Wash Buffer A (DUO82049), and ligation stock/ligase (DUO92014) was added for 30 minutes at 37°C. Slips were washed with Wash Buffer A, and then amplification stock/polymerase (DUO92014) was added for one hour and forty minutes at 37°C. Slides were washed with Wash Buffer B, and then mounted on slides with Duolink In Situ DAPI Mounting Medium (DUO82040) from Sigma Aldrich. Fifteen minutes after mounting, images were acquired on the Nikon Eclipse TS2 microscope at 40X magnification. All images were captured with an equal exposure time to ensure fluorescence can be compared. Images were then manually thresholded with ImageJ, and raw fluorescent signals per cell were quantified. Experiments were repeated three individual times, and representative images are shown.

### Membrane Fractionation

Control (C1-A) or FLOT2 KO CRISPR (F4-2) HeLa cells were scraped in ice-cold PBS, and a fraction of these cells were lysed in NP-40 as whole-cell lysate, while the rest of the pellet were fractionated with the subcellular protein fractionation kit for cultured cells, according to the manufacturers protocol (ThermoFisher 78840).

### Cholesterol/Cholesterol Ester-Glo Assay

Control (C1-A) HeLa cells were treated with simvastatin for 24 or 48 hours, and then the total cholesterol was quantified using the Cholesterol/Cholesterol Ester-Glo Assay Kit by Promega (J3190), according to the manufacturers instructions.

### Anchorage-independent soft agar growth assay

Bottom layer agar was made to reach a final concentration of 0.8% agar (DF0812-17-9; Fisher Scientific), 0.2% peptone (DF0118-17-0; Fisher Scientific), 1X DMEM (SLM-202-B; Millipore) with 20% FBS, 1% glutamine and 1% pen-strep, 2 mL was layered on bottom of each well in a 6-well plate, and allowed to set at room temperature. The top layer was made to reach a final concentration of 0.4% agar, 0.1% peptone, 1X DMEM with 20% FBS, 1% glutamine and 1% pen-strep, along with 35,000 cells suspended per well. Upon setting, a feeder layer of 10% FBS DMEM with DMSO, erlotinib, EGF or EGF + erlotinib was added. Plates were incubated at 37°C for three weeks, and colonies were counted using Nikon Eclipse TS2 microscope. Experiments were repeated three independent times.

### Quantitative reverse transcription PCR

Primers for EGFR ligands were constructed and cycled as follows and as previously described (52): **EPGN:** Forward-ATTCAACGCAATGACAGCACT and Reverse-TCCAGTTACCTTGCTGGGC. **EGF:** Forward-GACTTGGGAGCCTGAGCAGAA and Reverse-CATGCACAAGCGTGACTGGAGGT. **HB-EGF:** Forward-GGTGGTGCTGAAGCTCTTTC and Reverse-CCCCTTGCCTTTCTTCTTTC. **AREG:** Forward-GCCTCAGGCCATTATGC and Reverse-ACCTGTTCAACTCTGACTGA. **BTC:** Forward-TCTAGGTGCCCCAAGC and Reverse-GTGCAGACACCGATGA. **EREG:** Forward-AAAGTGTAGCTCTGACAT and Reverse-CTGTACCATCTGCAGAAATA. **TGFA:** Forward GCCCGCCCGTAAAATGGTCCCCTC and Reverse-GTCCACCTGGCCAAACTCCTCCTCTGGG. RNA was extracted with Trizol (Cat # 15596026; Thermo Fisher) per the manufacturer’s instructions from HeLa and H441 cells. Other reagents were used as previously described (53), and GAPDH served as loading control.

### Cell proliferation assays

35,000 HeLa or H441 cells were plated in triplicate in 12-well plates, per treatment condition. The following day (Day 0), cells were tryspinized and counted with ViaStain AOPI (acridine orange and propidium iodide) staining solution (CS2-0106) using a Cellometer K2 from Nexcelom Bioscience (Lawrence, MA). Other wells were washed with PBS and 1% FBS media was added to each well, along with corresponding drug or ligand treatments. Cells were trypsinized and counted with AOPI on Days 3, 5 and 7. Live cell counts were normalized to their corresponding Day 0 count to track growth over time. Experiments were repeated at least three independent times. Data were presented as an average ± SEM.

### Mouse models

Cells were counted with AOPI stain, and the appropriate numbers of live cells were suspended in PBS (nude mice injection) or a 1:1 ratio of PBS and Phenol Red-Free LDEV-Free Matrigel (356237; Corning) for NSG mice injection. Athymic nu/nu (Envigo) and NSG mice (Jackson labs) were housed and observed according to approved NCI-ACUC guidelines. Body weights were taken once weekly for 5–10 weeks or as required by humane endpoints for all mice. Subcutaneous (flank) injections were performed into 6– 8-week-old athymic nu/nu and NSG females in groups of 6 and 10 respectively. Subcutaneous tumor caliper measurements were taken once weekly in mice exhibiting palpable tumors until 5–10 weeks or humane endpoints. Tumor volumes were calculated according to the formula V = ½ (length × width^2^).

### Genomic Analysis

TCGA FPKM-UQ normalized mRNA-expression data downloaded and FLOT2 data extracted from NCI Genomics Data Commons (54) and the clinical data downloaded from cBioPortal (55) for each cancer type. Samples were grouped by the expression quartiles and the top and bottom quartile. Disease Free Survival were analyzed with the Kaplan-Meier method and groups compared with the log-rank test. Survival (https://cran.r-project.org/web/packages/survival/index.html) and Survminer (https://cran.r-project.org/web/packages/survminer/index.html) packages were used to generate the Kaplan-Meir plots. All statistical analysis performed with R (version 4.0.3) (56).

### Statistics

Comparisons were done using Student’s *t*-test with 2 tailed comparisons assuming equal variance or one-way ANOVA as indicated. A p-value of ≤0.05 was considered significant.

## Data and materials availability

The mass spectrometry proteomics data have been deposited to the ProteomeXchange Consortium via the PRIDE [1] partner repository with the dataset identifier PXD031988.

## This article contains supporting information

### Supporting Figures

Figure S1. In-Cell ELISA to assess effects of siRNA mediated knockdown of target genes on EGFR phosphorylation

Figure S2. Knockdown or knockout of FLOT2 increases EGFR phosphorylation, reduces basal EGFR and FLOT1 levels.

Figure S3. The genetic analysis of FLOT2 KO CRISPR clones, and co-Immunoprecipitation of FLOT2 and Cbl decreases with EGF stimulation.

Figure S4. Phosphorylation of multiple tyrosines on EGFR is increased upon FLOT2 KD. FLOT2 expression varies by cell line, and PC9 cells are sensitive to erlotinib.

Figure S5. FLOT2 KD induces MAPK activation dependent on EGFR signaling, but not MET, FAK, VEGFR, KIT, AXL, RET signaling.

Figure S6. EGF induces EGFR dimerization as visualized by proximity ligation assay. FLOT2 KO reduces EGFR in the membrane fraction, and EGFR downregulation by FLOT2 loss is not rescued by cholesterol inhibition.

Figure S7. Trafficking of EGFR to the lysosome in FLOT2 KO cells.

Figure S8. FLOT2 interacts with EGFR.

Figure S9. HeLa growth in presence of erlotinib and after KD by multiple FLOT2 siRNAs.

Figure S10. FLOT2 expression and disease free survival varies depending on cancer subtype.

### Supporting Tables

Table S1. All proteins identified with SILAC mass spectrometry in the complex with Cbl without treatment (L) and with 30 min EGF stimulation (H).

Table S2. EGFR ligand expression in HeLa and H441 cells.

## Acknowledgments

We thank all members of the Lipkowitz lab in the Women’s Malignancies Branch, Center for Cancer Research, National Cancer Institute for their support and review of this manuscript.

## Funding

This research was supported by the Intramural Research Program of the National Cancer Institute (ZIA BC 010977).

## Conflict of interest

The authors declare that they have no conflicts of interest with the contents of this article.

## Abbreviations and Nomenclature

Cbl: Casitas B-lineage lymphoma proto-oncogene
EGF: epidermal growth factor
EGFR: epidermal growth factor receptor
Ub: ubiquitination
a.u: arbitrary units
FLOT1: Flotillin 1
FLOT2: Flotillin 2
KD: knockdown
KO: knockout
RF: ring finger
RTK: receptor tyrosine kinase
SILAC: Stable Isotope Labeling with Amino Acids in Cell Culture
WB: Western blotting
PLA: proximity ligation assay
AOPI: acridine orange and propidium iodide
WCL: whole-cell lysates
IP: immunoprecipitation

## Notes

### Competing Interest Statement

The authors have declared no competing interest.

### Summary of Updates

Table 1 has been created to create clarity, main figures have been updated with new data to improve the strength of the manuscript. Supplemental figures have been updated to improve the strength of the manuscript

