## Supporting Information for "Flotillin-2 regulates EGFR activation, degradation, and cancer growth"

**Contents:**

**Supporting Figures S1-S10**

**Supporting Table S1 and S2 Legend**

**Supporting Figures:**

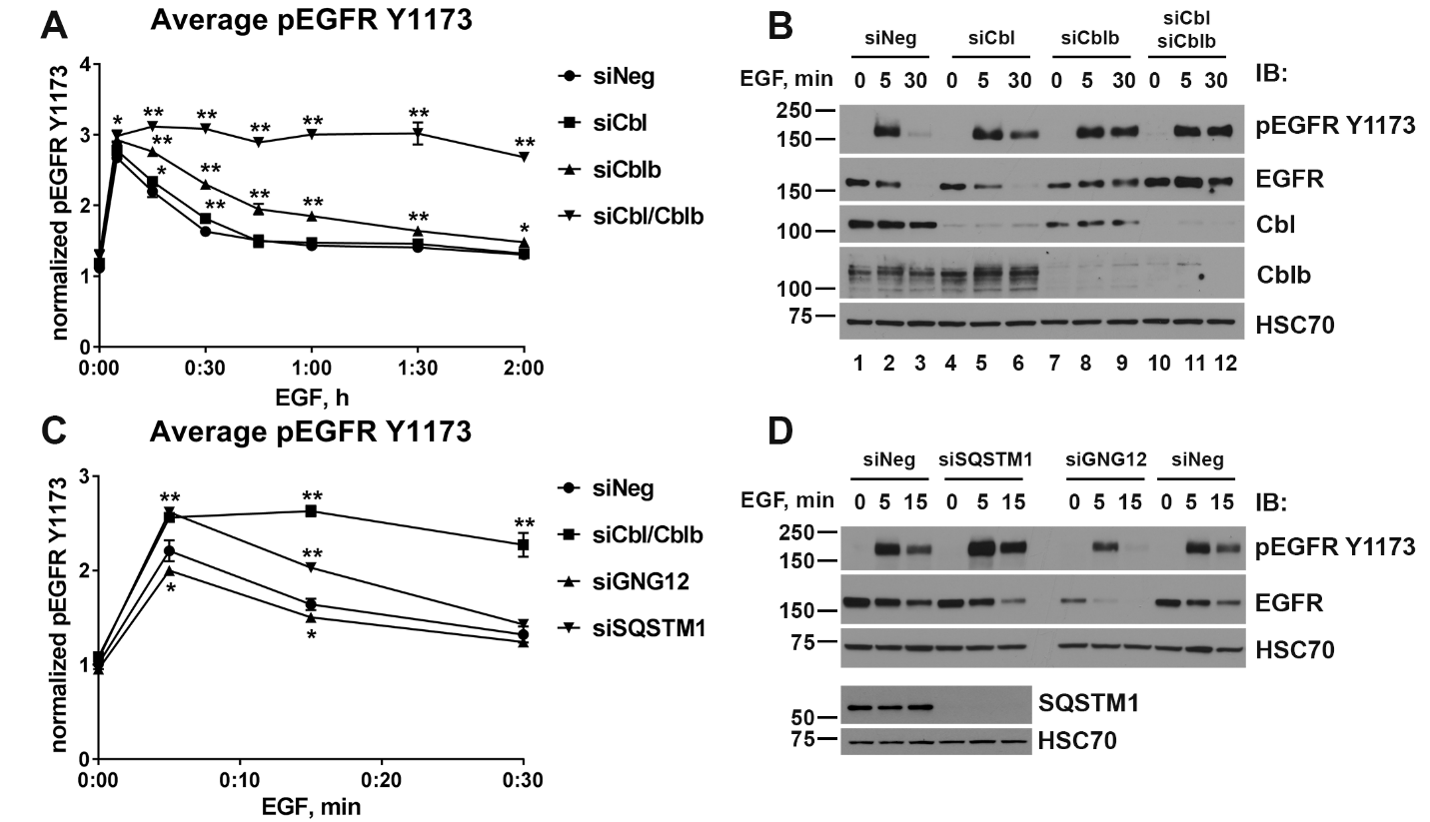

**Figure S1. In-Cell ELISA to assess effects of siRNA mediated knockdown of target genes on EGFR phosphorylation. (A)** HeLa cells were transfected with non-targeting siRNA (siNeg, negative control), siCbl, siCblb, and siCbl/Cblb and seeded on 96-well plate. Phospho-EGFR ELISA was used to detect the levels of EGFR phosphorylated at Y1173 upon EGF stimulation for indicated time periods. The pEGFR Y1173 signal was normalized to the Janus-green whole-cell staining signal and the average values (± SD) were plotted versus EGF stimulation time. (B) Western blot analysis of the same transfection as in A demonstrates the efficient knockdown of Cbl and Cblb as well as delay in EGFR degradation upon stimulation with 100 ng/mL EGF for 5 and 30 minutes in Cbl/Cblb KD cells. (C) HeLa cells were transfected with siRNAs targeting siNeg (negative control), Cbl/Cblb (positive control), GNG12 and SQSTM1 and analyzed as in A. (D) HeLa cells were transfected with siNeg, siSQSTM1 and siGNG12, stimulated with 100 ng/mL EGF for 5 or 15 min and subjected to the western blot with the indicated antibodies. In A and C, asterisk (*) denotes p < 0.05, (**) indicates p < 0.01 as compared to the corresponding time points for siNeg using Student’s *t*-test. MW in kDa is shown to the left of the panels in B and D.

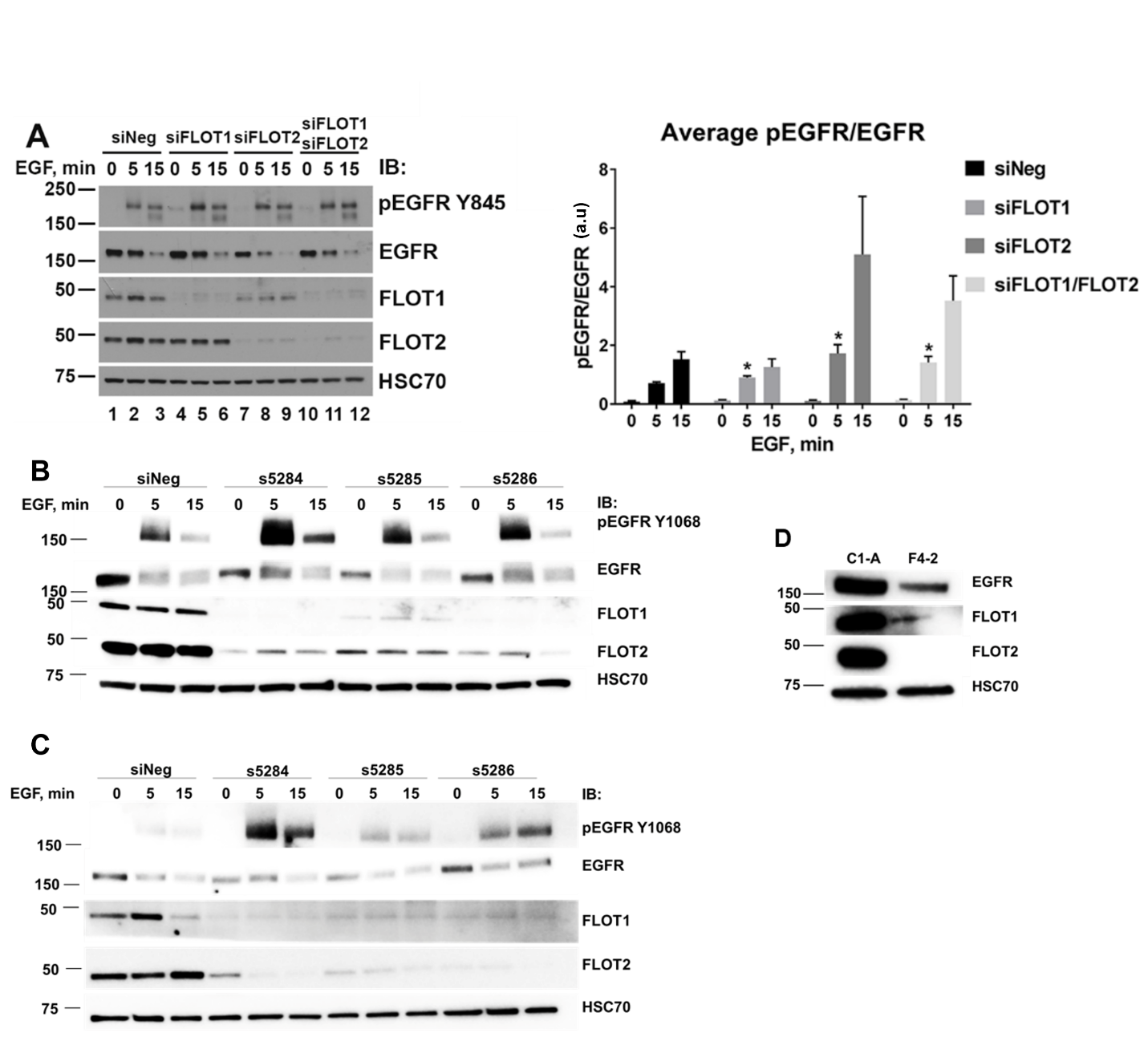

**Figure S2. Knockdown or knockout of FLOT2 increases EGFR phosphorylation, reduces basal EGFR and FLOT1 levels. (A)** HeLa cells were transfected for 48 hours with control siRNA (siNeg) or siRNAs targeting FLOT1 and FLOT2 separately or together. The lysates were analyzed by western blot to detect pEGFR Y845, EGFR, FLOT1, FLOT2 and HSC70. EGFR phosphorylation at Y845 upon EGF stimulation in HeLa cells is shown as an average ratio of pEGFR/EGFR ± SEM calculated by densitometry analysis of western blot in A for four independent experiments, with a.u indicating arbitrary units. Asterisk (*) denotes p < 0.05 as compared to the corresponding time points for siNeg using Student’s *t*-test. HeLa (B) or H441 (C) were transfected with the indicated siRNA for 48 or 72 hours, respectively, then serum starved for 3 hours and stimulated with 25 ng/mL EGF for the indicated time points. Lysates were analyzed by western blot to detect pEGFR (Y1068), EGFR, FLOT1, FLOT2 and HSC70 (loading control). (D) HeLa control (C1-A) or FLOT2 (F4-2) CRISPR clone lysates were analyzed by western blot for EGFR, FLOT1, FLOT2 and HSC70 (loading control).

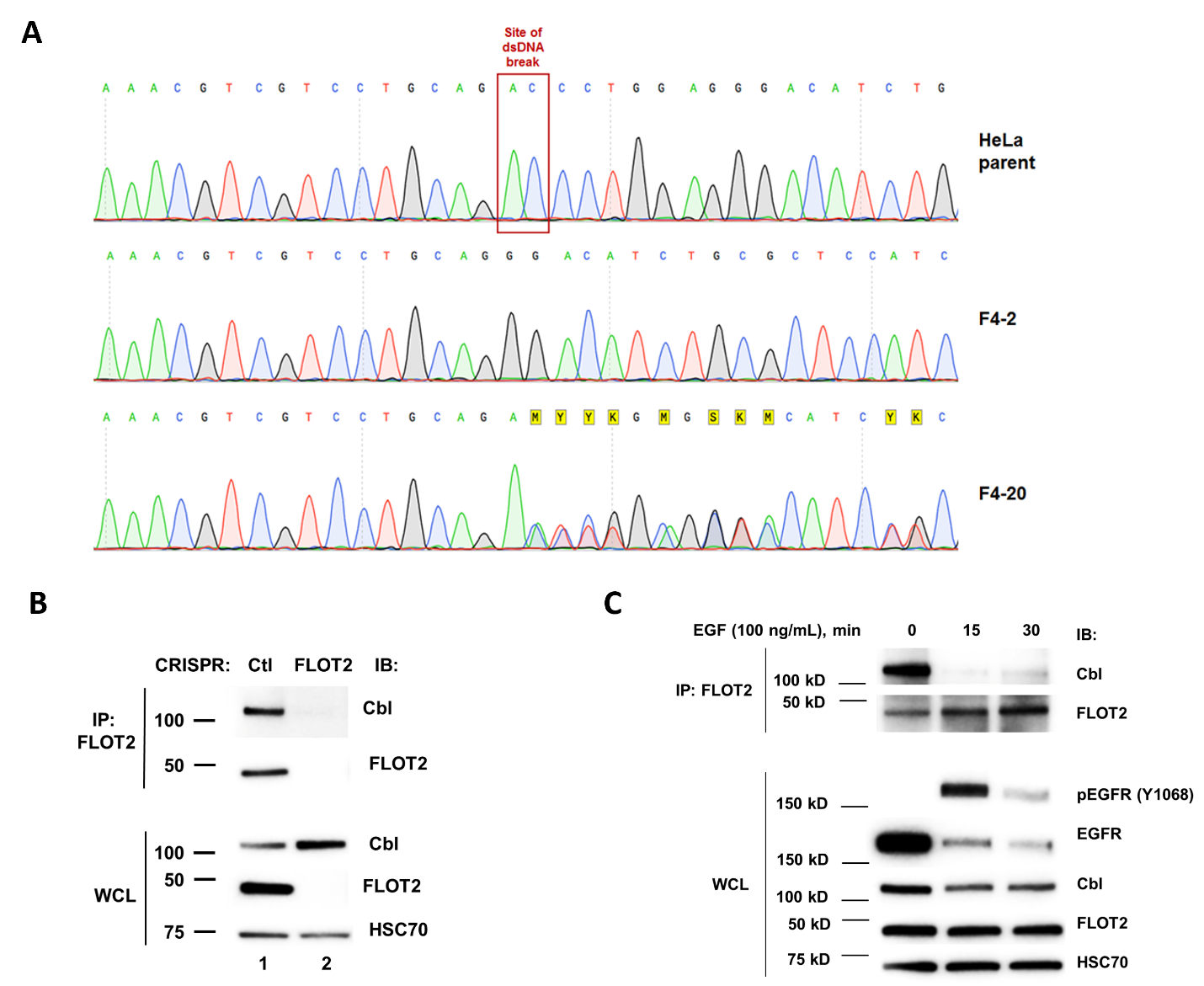

**Figure S3. The genetic analysis of FLOT2 KO CRISPR clones, and co-Immunoprecipitation of FLOT2 and Cbl decreases with EGF stimulation.**

**(A)** Genetic sequencing analysis was performed to compare the genomic sequence of FLOT2 KO clones F4-2 and F4-20 to parent HeLa cells. CRISPR Clone F4-2 has a homozygous deletion of 9 bases leading to an in-frame deletion of 3 amino acids. This resulted in complete loss of the protein. Clone F4-20 showed evidence for two frame shift mutations and again resulted in complete loss of FLOT2 protein. (B) HeLa CRISPR control (C1-A) or FLOT2 KO (F4-2) lysates were immunoprecipitated with FLOT2 antibody, and then probed on a western blot for Cbl and FLOT2, with whole-cell lysates being probed for Cbl, FLOT2 and HSC70 (loading control). (C) HeLa cells were serum starved for 3 hours and stimulated with 100 ng/mL EGF for the indicated time points, lysed, and immunoprecipitated with FLOT2 antibody. FLOT2 IP was probed on a western blot for Cbl and FLOT2, and whole-cell lysates were probed for pEGFR (Y1068), EGFR, Cbl, FLOT2 and HSC70 (loading control).

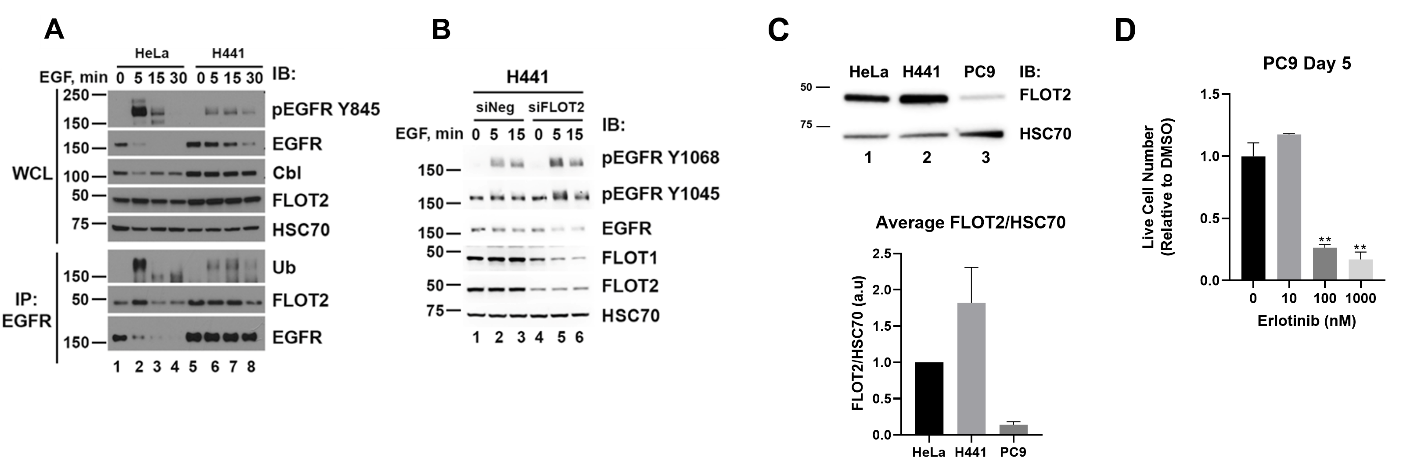

**Figure S4. Phosphorylation of multiple tyrosines on EGFR is increased upon FLOT2 KD. FLOT2 expression varies by cell line, and PC9 cells are sensitive to erlotinib.**

**(A)** HeLa and H441 cells were treated with 100 ng/mL EGF for indicated times and the levels of pEGFR Y845, EGFR, Cbl, FLOT2 and HSC70 were compared between the cell lines by western blot of whole cell lysates (WCL). EGFR was immunoprecipitated and probed for ubiquitin (Ub), FLOT2 and EGFR. (B) H441 cells were transfected with siNeg and siFLOT2 for 72 hours and the lysates were analyzed by western blot with the indicated antibodies. MW in kDa is shown to the left of the western blot panels. (C) HeLa, H441 and PC9 cells were lysed and probed on a western blot for FLOT2 and HSC70 (loading control). Average EGFR/HSC70 ± SEM was calculated by densitometry analysis of western blot for three independent experiments, with a.u indicating arbitrary units. MW in kDa is shown to the left of the western blot panels. (D) PC9 cells were plated, treated for five days with increasing concentrations of erlotinib, and live cells were counted by AOPI staining. Asterisk (**) indicates p < 0.01 as compared to DMSO control using Student’s *t*-test.

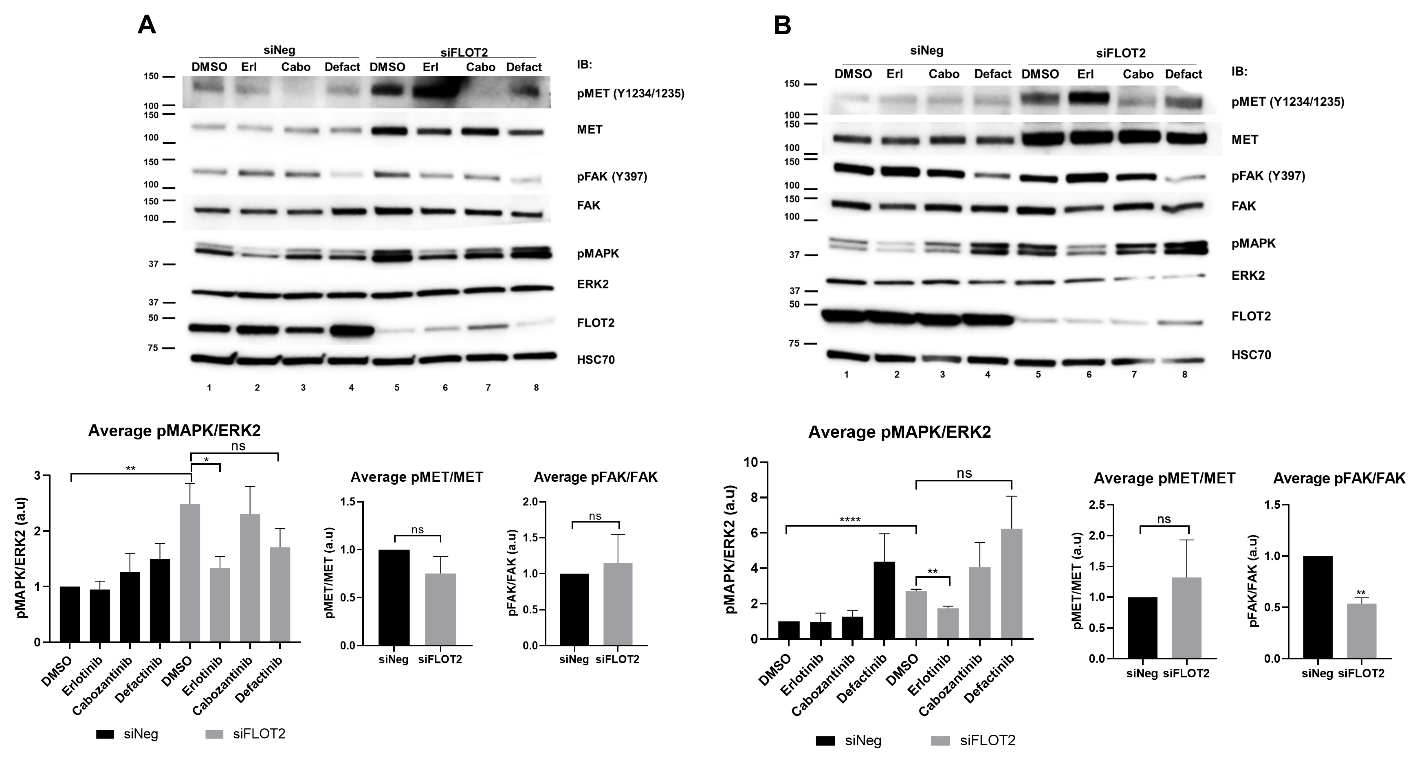
**Figure S5. FLOT2 KD induces MAPK activation dependent on EGFR signaling, but not MET, FAK, VEGFR, KIT, AXL, RET signaling.**

**(A)**	H441 cells were transfected with siNeg or siFLOT2 for 72 hours, and then treated with DMSO, erlotinib (10 µM), cabozantinib (1 µM), or defactinib (1 µM) in 1% FBS RPMI for 24 hours. Average pMAPK/ERK2 ± SEM, pMET/MET ± SEM and pFAK/FAK ± SEM were calculated by densitometry analysis of western blot in four independent experiments, with a.u indicating arbitrary units. Asterisk (**) indicates p < 0.01 and (*) indicates p<0.05 using Student’s *t*-test. MW in kDa is shown to the left of the western blot panels. (B) PC9 cells were transfected with siNeg or siFLOT2 for 48 hours, and then treated with DMSO, erlotinib (10 nM), cabozantinib (1 µM), or defactinib (10 µM) in 1% FBS RPMI for 24 hours. Average pMAPK/ERK2 ± SEM, pMET/MET ± SEM and pFAK/FAK ± SEM were calculated by densitometry analysis of western blot in three independent experiments, with a.u indicating arbitrary units. Asterisk (**) indicates p < 0.01 and (****) indicates p<0.0001 using Student’s *t*-test. MW in kDa is shown to the left of the western blot panels.

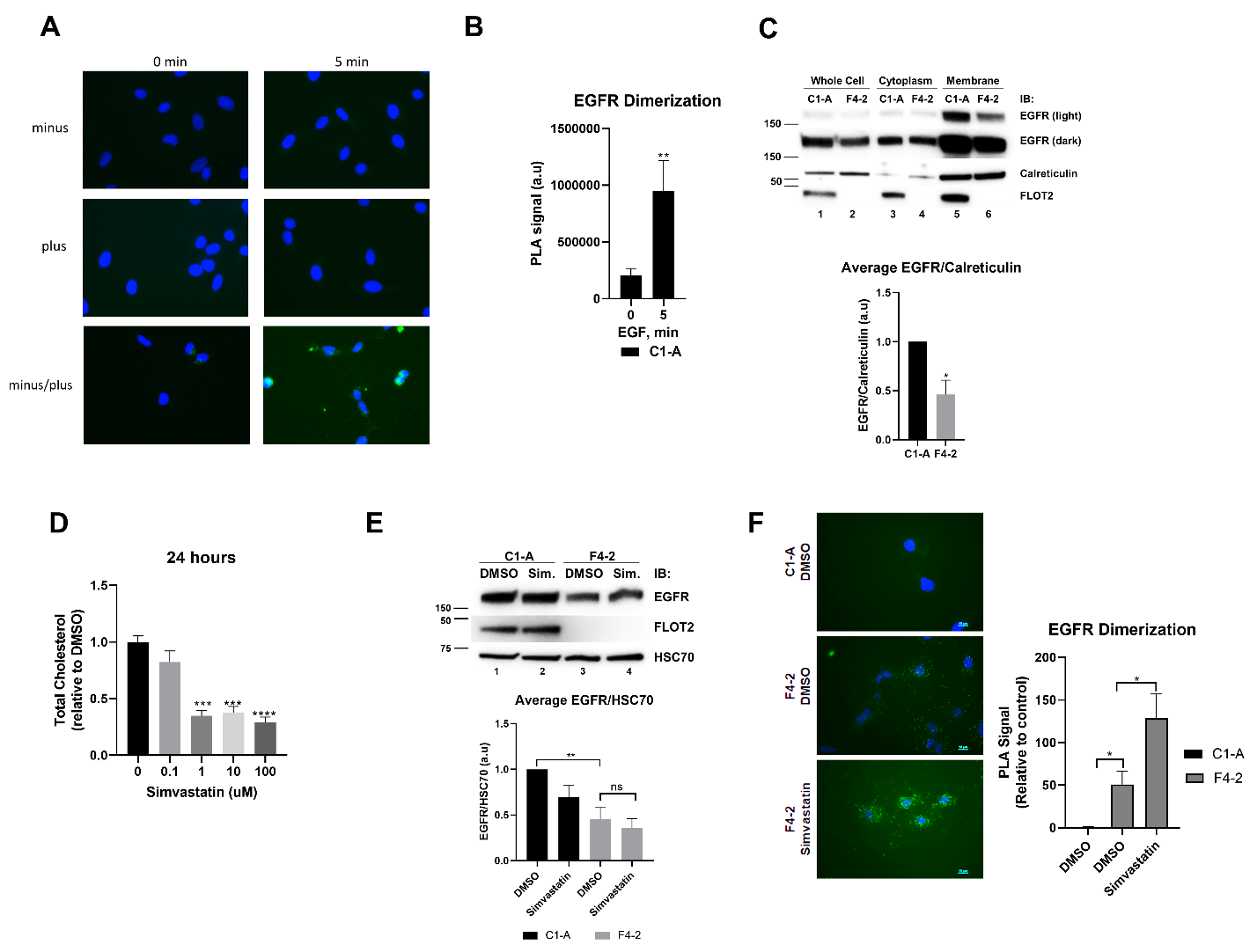
**Figure S6. EGF induces EGFR dimerization as visualized by proximity ligation assay. FLOT2 KO reduces EGFR in the membrane fraction, and EGFR downregulation by FLOT2 loss is not rescued by cholesterol inhibition. (A)** Control (C1-A) HeLa cells were stimulated with EGF for 5 minutes and subsequently treated with PLUS alone, MINUS alone, or PLUS/MINUS combined and counterstained with DAPI. No detectable green fluorescence under high exposure indicates no background or non-specific signal. (B) Control (C1-A) HeLa cells were stimulated with EGF for 5 minutes and subsequently treated with PLUS and MINUS probes conjugated to EGFR ABs in a PLA. Dimerized EGFR was visualized by green fluorescence, when both PLUS and MINUS probes bound to EGFR were within close proximity. Fluorescence was quantified using ImageJ, and the average signal in arbitrary units (a.u) (± SEM) of three independent experiments were plotted. Asterisks (**) denotes p < 0.01 using Students *t*-test. (C) Control (C1-A) or FLOT2 KO CRISPR (F4-2) HeLa cells were collected and subjected to subcellular fractionation, resulting in whole-cell lysate, cytoplasmic, and membrane associated proteins. Lysates were probed by western blot for EGFR, calreticulin and FLOT2. Average EGFR/Calreticulin ± SEM was calculated by densitometry analysis of western blot in three independent experiments, with a.u indicating arbitrary units. Asterisks (*) denotes p < 0.05 using Students *t*-test. MW in kDa is shown to the left of the western blot panels. (D) Control (C1-A) HeLa cells were treated with increasing concentrations of simvastatin for 24 hours and total cholesterol was measured using a Cholesterol-Glo Assay. Asterisks (**) denotes p <0.01, (***) denotes p <0.001, (****) denotes p <0.0001 using Student’s *t*-test. (E) Control (C1-A) or FLOT2 KO CRISPR (F4-2) HeLa cells were treated with 1 µM simvastatin for 24 hours, and lysates were subjected to western blot analysis of EGFR, FLOT2 and HSC70 (loading control). Average EGFR/HSC70 ± SEM was calculated by densitometry analysis of western blot in five independent experiments, with a.u indicating arbitrary units. Asterisks (**) denotes p <0.01 and ns indicates p>0.05 using Student’s *t*-test. MW in kDa is shown to the left of the western blot panels. (F) Control (C1-A) HeLa cells were treated with DMSO control, and FLOT2 KO CRISPR (F4-2) HeLa cells were treated with DMSO control or 1 µM simvastatin in 1% FBS RPMI for 24 hours, and subsequently treated with PLUS alone, MINUS alone, or PLUS/MINUS combined and counterstained with DAPI. Asterisks (*) denotes p < 0.05, (***) denotes p <0.001 using Students *t*-test.

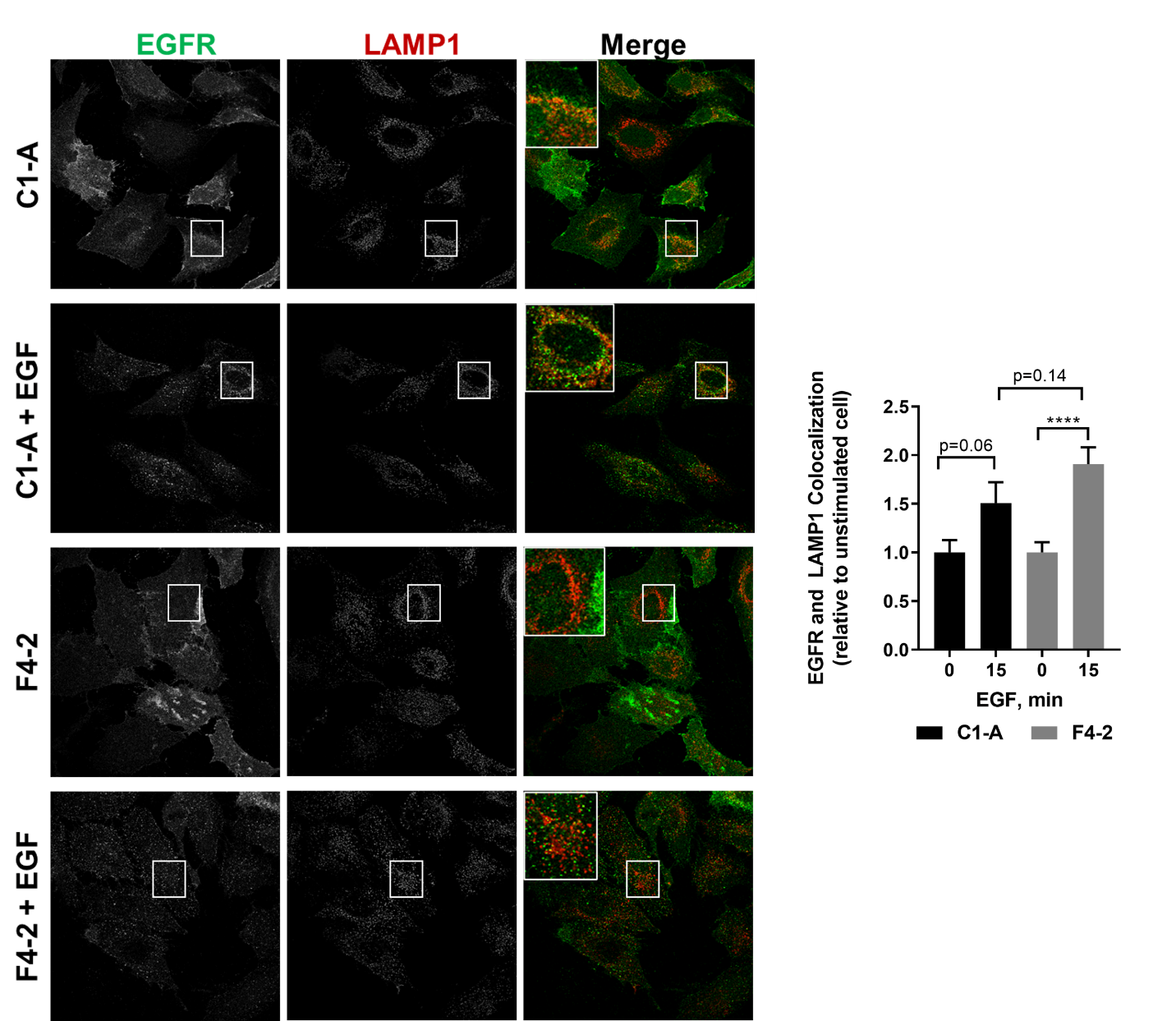
**Figure S7. Trafficking of EGFR to the lysosome in FLOT2 KO cells.**

Control (C1-A) and FLOT2 (F4-2) CRISPR KO HeLa cells were treated with or without 25 ng/mL EGF for 15 minutes, and were subsequently stained with EGFR AB and LAMP1 AB, followed by Alexa-Fluor 488 and Alexa-Fluor 594 fluorescent secondary AB, respectively. Overlapping EGFR and LAMP1 pixels were calculated using ImageJ. Stimulated C1-A cells were normalized to unstimulated C1-A cells, and stimulated F4-2 cells were normalized to unstimulated F4-2 cells. The graph shows the average (± SEM) of three independent experiments. Asterisks (****) denotes p < 0.0001 using Student’s *t*-test. p values >0.05 are indicated on the graph.

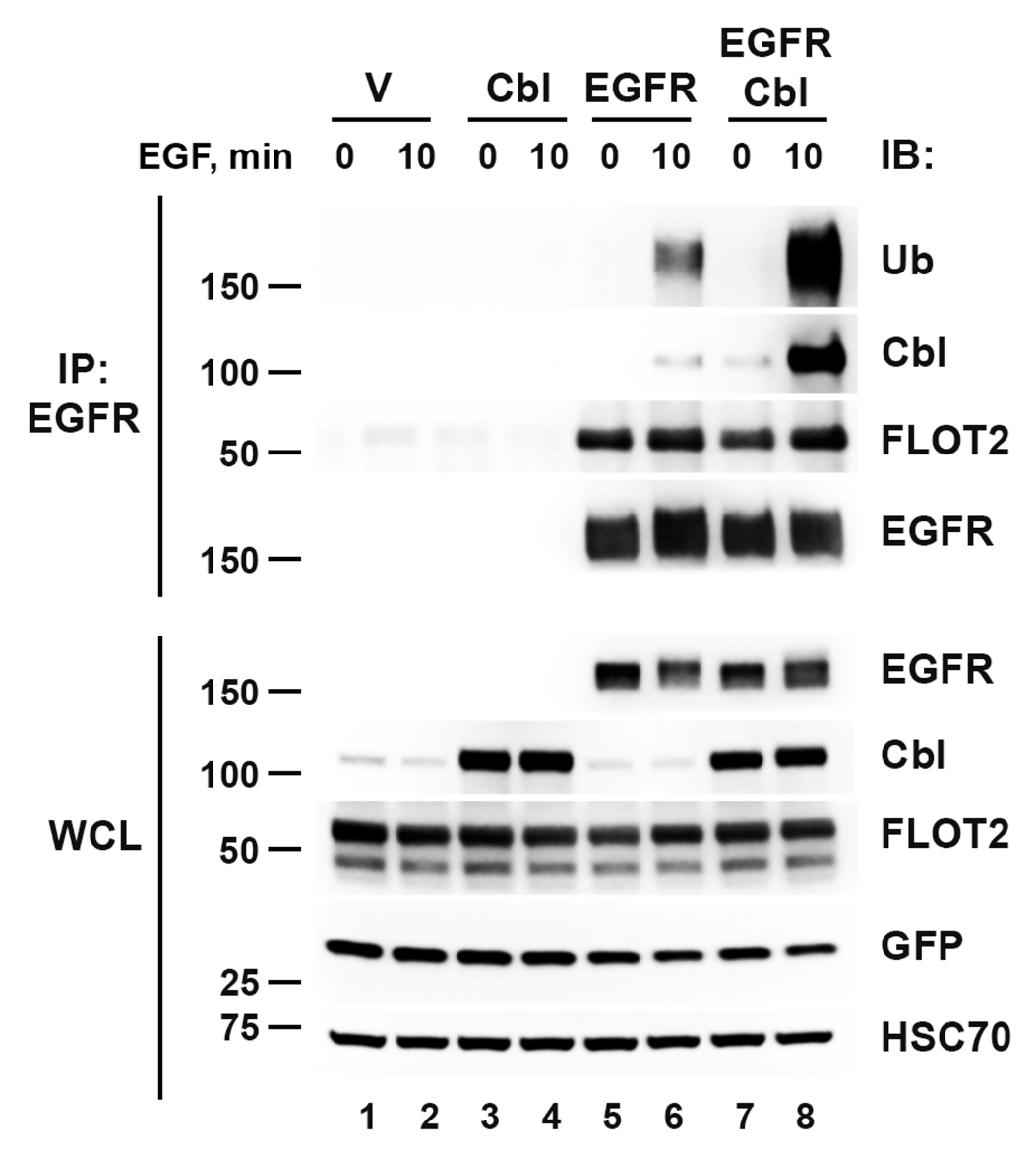

**Figure S8.** **FLOT2 interacts with EGFR.** HEK293T cells were transfected with WT FLAG-FLOT2 in combination with empty Vector (V, control), Cbl, EGFR or Cbl/EGFR as indicated. GFP plasmid was used as a transfection control. 48-hours post transfection the cells were stimulated with 100 ng/mL EGF for 10 min and the lysates were analyzed by western blot for EGFR, Cbl, FLOT2, GFP and HSC70. EGFR was immunoprecipitated from the lysates and analyzed by western blot with the indicated antibodies. Overexpressed FLOT2 was detected either with anti-FLOT2 antibodies. MW in kDa is shown to the left of the western blot panels.

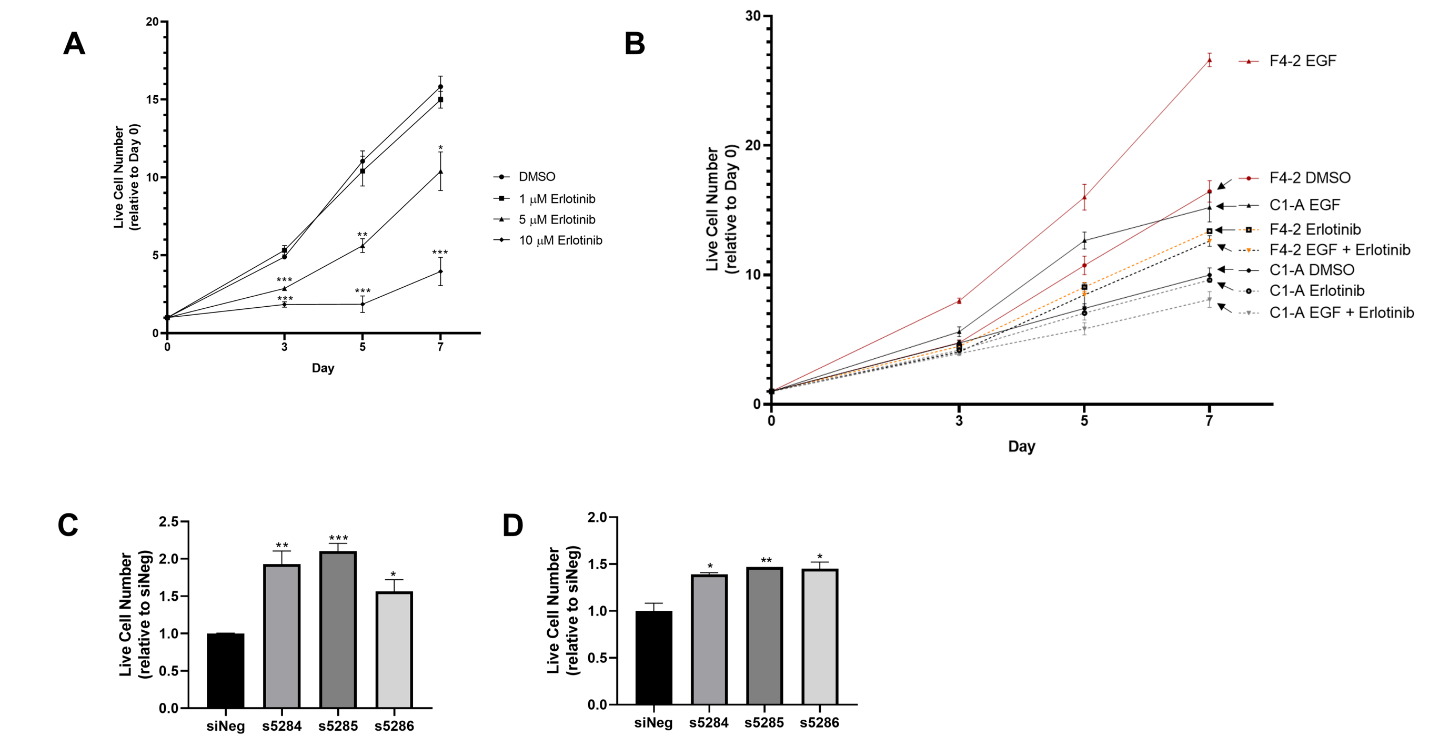
**Figure S9. HeLa growth in presence of erlotinib and after KD by multiple FLOT2 siRNAs.(A)** HeLa parent cells were plated and treated next day with DMSO (control), 1 µM erlotinib, 5 µM erlotinib or 10 µM erlotinib, and subsequently counted with AOPI staining for viable cells on Day 0 (day of treatment), 3, 5 and 7. Cell number was normalized to the Day 0 counts. The graph shows the average (± SEM) of three independent experiments. Asterisks (*) denotes p <0.05, (**) denotes p <0.01, and (***) denotes p <0.001 using students *t*-test. (B) Control (C1-A) and FLOT2 KO CRISPR (F4-2) HeLa cells were plated and treated with DMSO (control), 1 µM erlotinib, 25 ng/mL EGF, or erlotinib + EGF, and subsequently counted with AOPI staining for viable cells on Day 0, 3, 5 and 7. Cell number was normalized to the Day 0 counts of each cell line. HeLa (C) or H441 (D) cells were transfected with indicated siRNA for 48 hours, replated, and subsequently counted with AOPI staining for viable cells on Day 0 (day following re-plating) and Day 7. Cell number was normalized to the Day 0 counts, and plotted the average of three independent experiments. Asterisks (*) denotes p <0.05, (**) denotes p <0.01, and (***) denotes p <0.001 using One-way ANOVA testing with Dunnett multiple comparisons post-test.

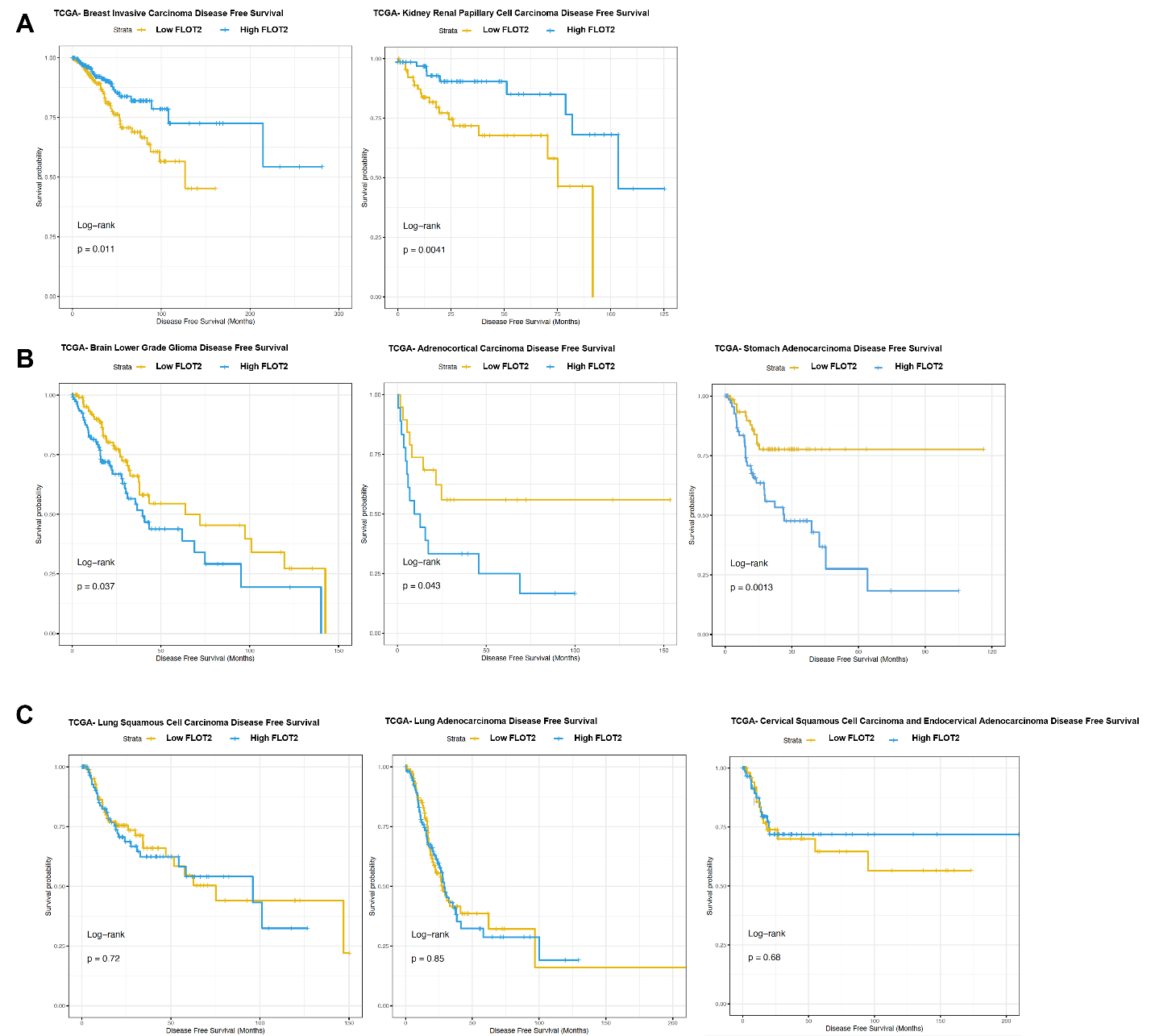
**Figure S10. FLOT2 expression and disease free survival varies depending on cancer subtype.**

**(A)** TCGA data which predicts high FLOT2 expression improves disease free survival in a statistically significant manner. (B) TCGA data which predicts low FLOT2 expression improves disease free survival in a statistically significant manner. (C) TCGA data which predicts FLOT2 expression has no statistical effect on disease free survival

**Table S1.** All proteins identified with SILAC mass spectrometry in the complex with Cbl without treatment (L) and with 30 min EGF stimulation (H). The data for 5 independent experiments is shown. In experiments 1 and 2 mixed IP Cbl was directly used to generate peptides for MS. For experiments 3, 4 and 5, mixed IP Cbl was separated by SDS-PAGE, divided into fractions and analyzed with MS.

**Table S2. EGFR ligand expression in HeLa and H441 cells.** In three independent experiments, RT-qPCR was performed to determine delta Ct values (Target Gene- GAPDH Gene Expression). N/D indicates no detectable mRNA. Two-tailed students *t-*test was performed to determine statistical significance between HeLa and H441. N/T indicates not tested.

| Gene | H441 ΔCt | HeLa ΔCt | p value |
| --- | --- | --- | --- |
| EPGN | 13.93694 | 13.62133 | 0.76 |
| EGF | 13.40834 | 13.33473 | 0.895 |
| HB-EGF | 10.23345 | 13.04033 | 0.128 |
| AREG | 2.841667 | 11.75933 | 0.000789 |
| BTC | 13.47 | N/D | N/T |
| TGFA | 10.80367 | 17.095 | 0.0558 |
| EREG | 12.785 | 16.165 | 0.13 |
